# The small genome size ensures adaptive flexibility for an alpine ginger

**DOI:** 10.1101/2024.12.31.630872

**Authors:** Qing-Song Xiao, Tomáš Fér, Wen Guo, Hong-Fan Chen, Li Li, Jian-Li Zhao

**Author notes:** **Corresponding author.** Ministry of Education Key Laboratory for Transboundary Ecosecurity of Southwest China, Yunnan Key Laboratory of Plant Reproductive Adaptation and Evolutionary Ecology, Institute of Biodiversity, School of Ecology and Environmental Science, Yunnan University, Kunming, 650504, Yunnan, China *E-mail addresses:* (J. L. Zhao).

## Abstract

Understanding the proximate and ultimate causes of genome size (GS) variation is the focus of much research. However, the extent and causes of intraspecific variation in GS is debated and poorly understood. This study aims to test the large-genome constraint hypothesis through the variations of intraspecific GS. GS was measured in 53 *Roscoea tibetica* populations from the Hengduan Mountains using flow cytometry. Stomatal size and density were collected from the wild populations and common garden populations. Associations between GS and environmental factors, stomatal traits were explored. We found that high GS variability was positively correlated with most environmental factors but negatively correlated with solar radiation during the growing season. The environment, rather than geography, significantly influenced the variations in GS. The stomatal traits measured in the wild were significantly correlated with GS, but no such correlations were detected in the common garden. Populations in the common garden have larger stomatal size and lower stomatal density. Populations with smaller GS present larger degree of stomatal traits variation from wild to common garden. Our findings suggests that intraspecific GS has experienced adaptive evolution driven by environmental stress and the evolution of intraspecific GS can be explained by the large-genome constraint hypothesis. Smaller GS is more advantageous to the alpine ginger to adapt to alpine habitat and thrive in changing habitat.

## 1. Introduction

Variations in genome size (GS) play a crucial role in evolution and ecological adaptation (Beaulieu et al., 2008; Bennett, 1987; Bureš et al., 2024; Marais et al., 2020; Schubert and Vu, 2016; Yang et al., 2013). Although numerous studies have explored the reasons for GS variation, why GS varies substantially in living beings remains a fascinating but puzzling question (Armstrong et al., 2023; Dkennedy and Norman, 2005; Sanders et al., 2021).

Several hypotheses have been proposed from different perspectives regarding the variation of GS and the origin of genome complexity. The main theories assume that natural selection is the primary factor leading to the evolution of GS. Examples of these theories include the nucleotypic (Bennett, 1977; Gregory and Hebert, 1999), nucleoskeletal (Cavalier-Smith, 1978, 2005), genome-streamlining (Hessen et al., 2010) and large-genome-constraint hypotheses (Bureš et al., 2024; Knight et al., 2005). The nucleotypic and nucleoskeletal hypotheses assumes a strong correlation between GS and nuclear/cell volumes. Accordingly, GS can affect the phenotypic characteristics of organisms, including individual size, development time, and cell size, among others. Natural selection acting on phenotypes, in turn, indirectly drives the evolution of GS (Beaulieu et al., 2008; Bhadra et al., 2023; Roddy et al., 2020; Schley et al., 2022; Simonin and Roddy, 2018; Theroux-Rancourt et al., 2021). The genome-streamlining hypothesis proposes that metabolic resources such as nitrogen (N) and phosphorus (P) play an important role in GS selection. As N and P are the main components of DNA, individuals with larger genomes are at a disadvantage when N and P are limited (Acquisti et al., 2009; Faizullah et al., 2021; Guignard et al., 2016; Hessen et al., 2010; Leitch et al., 2014). However, when N and P are sufficient, plants with larger genomes accumulate more aboveground biomass and exhibit greater competitiveness relative to small genomes (Peng et al., 2022).

The large-genome constraint hypothesis integrates ideas from these three hypotheses. It suggests that a larger GS produces a larger cell volume, which limits physiological activity (Šmarda et al., 2023; Theroux-Rancourt et al., 2021; Veselý et al., 2020), decreases the cell division rate (Šímová and Herben, 2012) and increases plant N and P requirements (Peng et al., 2022). The current consensus on the effects of GS on cell size is that GS determines the minimum cell size rather than cell size in general (Bennett, 1987; Bhadra et al., 2023). The nucleotypic, nucleoskeletal, and genome-streamlining hypotheses are not equivalent to the large-genome-constraint hypothesis. The large-genome-constraint hypothesis is an overarching insight offering a broader perspective: it implies that large genomes restrict the flexibility of species or individuals in genome-size-related traits, thereby narrowing the range of environmental conditions in which a species or individual can thrive. Thus, a larger GS may be disadvantageous for species in harsh environments.

However, the evidence for the large-genome-constraint hypothesis come primarily from between species comparisons and phenotypic variation in the wild (Knight et al., 2005; Peng et al., 2022; Šmarda et al., 2010; Šmarda et al., 2023; Theroux-Rancourt et al., 2021; Veselý et al., 2020). Whether the hypothesis is applicable to GS variability at the intraspecific level remains largely unknown. Additionally, how cultivation affects the correlation between GS variation and phenotypic characteristics has not been elucidated. The GS of an individual should be identical regardless of whether they are growing in the wild or in the common garden. Assuming that the large-genome constraint hypothesis about GS evolution is universal, the correlation between intraspecific GS and phenotypic variation, such as cell size, should also not be constant between populations in the wild and common gardens because the plant traits of smaller GS are assumed to present more flexible the plant traits of larger GS. Till date, this assumption has not been tested within species by comparing the phenotypic characteristics of wild and common gardens.

*Roscoea tibetica* (Zingiberaceae), is one of an alpine gingers in *Roscoea*, is widely distributed in the biodiversity hotspot area of the Hengduan Mountains (HDMs) from low-altitude forests to high-altitude meadows with large morphological variations (Fig. 1 a and b) (Li et al., 2021). A previous study has revealed that *R. tibetica* has three ecotypes based on morphological and evolutionary differences: alpine meadow type (AM), high-altitude forest type (HF) and low-altitude forest type (LF) (Li et al., 2023). Large morphological and ecological variations of *R. tibetica* provide us an opportunity to test the applicability of the large-genome constraint hypothesis to intraspecific GS variation. Here, according to the large-genome constraint hypothesis for *R. tibetica*, we assumed that 1) the harsher the environment in the higher elevation has smaller GS in the wild populations and 2) the smaller of the GS exhibit larger variation of morphological traits, such as stomatal size and density, between wild and common garden populations. To test these hypotheses, correlations between GS and environmental factors and between GS and morphological traits under wild and common garden conditions were evaluated. Based on these correlations, we aimed to determine whether the large-genome constraint hypothesis can sufficiently explain the geographical patterns of intraspecific GS variation.

**Fig. 1.**
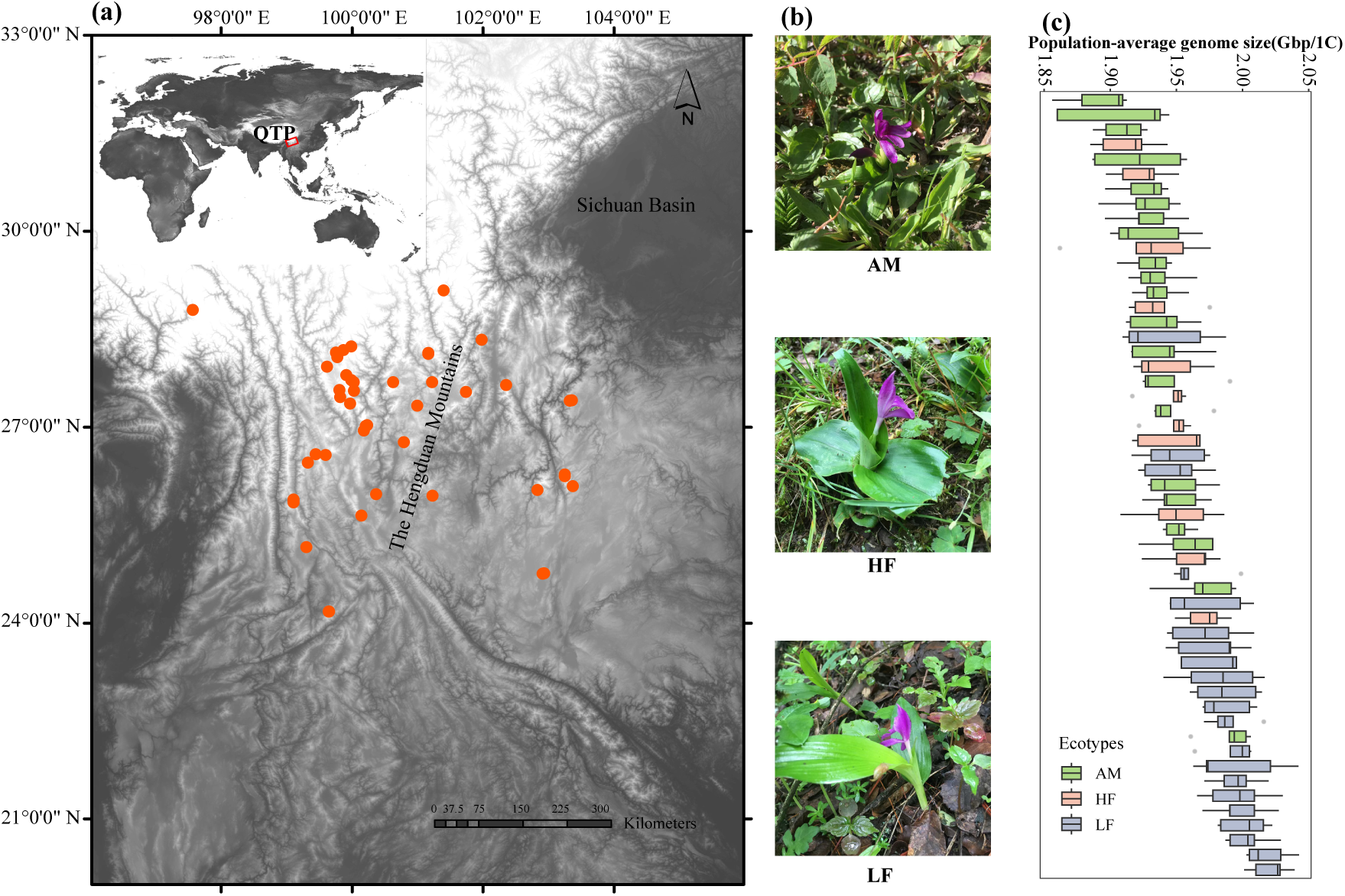
Sampling sites across the entire distribution range and intraspecific genome size (GS) variation of *R. tibetica*. (a) map of the collection locations of *R. tibetica*. The panorama at the upper left corner illustrates the distribution of *R. tibetica* in the southeast of Qinghai-Tibetan Plateau (QTP). (b) morphological diversity in *R. tibetica*. AM, alpine meadow type; HF, high-altitude forest type; LF is low-altitude forest type. (c) the distribution of population-mean GS for 53 wild populations of *R. tibetica*. Populations are arranged by mean GS values. The whiskers represent the standard deviation, and the grey dots represent the outliers.

## 2. Material and methods

### 2.1 Sampling

We conducted an extensive sampling of *R. tibetica* across its distribution range from 2400-3800 m in altitude. To compare the variations of morphological traits, such as cell size and density, we also collect samples from common garden. Per population, approximately 20 individual living plants with rhizomes were collected for common garden cultivation at Yunnan University, Kunming, China (102.8548° E, 24.8291° N). Young, fresh leaves were collected from 53 common garden populations for GS measurements, including 21 LF, 11 HF, and 21 AM populations (Fig. 1 a; Table S1, S2). Since collecting mature leaves for corresponding 53 populations was very challenging during the flowering stage in one year, we tried our best to collect mature leaves for measurements from 22 wild populations and 17 common garden populations (Table S3) to avoid annual fluctuations of morphological traits from confounding the results.

### 2.2 GS measurements

The nuclear DNA content (1C) of *R. tibetica* was measured by flow cytometry (FCM). Using internal standard method, the nucleus of the samples was stained using a two-step method with propidium iodide (PI) fluorescent dye (Greilhuber et al., 2007; Pellicer et al., 2021). Leaves of five individuals were randomly selected from each population for GS measurements, and in total, the GS of 265 individuals were measured. We mixed 50 mg of leaves of *R. tibetica* and 50 mg internal standard tomato (1C = 0.958 Gb) with 500 µL nuclei extraction buffer in a plastic Petri dish. The leaf tissues were chopped to dissociate the cells into a buffer (CyStain^TM^, Sysmex Partc GmbH, Czech) and incubated for 30–90 s at room temperature (∼25°C). Cell solutions were filtered through 50-µm CellTris^TM^ filter into centrifugal tubes with 2 mL PI staining solution (CyStain^TM^), and incubated for 15–30 min in the dark at room temperature. Finally, the prepared sample solutions were analyzed using a flow cytometer (CyFlow Space, Sysmex Europe SE, Norderstedt, Germany). The minimum number of nuclei for the sample to be tested was set to 5000 nuclei, and the coefficients of variation (CV) of the sample and standard peaks in the flow histogram were always less than 5%, with a mean of 3.8% (Table S2). The 1C DNA content of the samples was calculated using the following equation: nuclear DNA content (Gb) = 0.958 × (mean position of the reference standard peak /mean position of the reference sample peak) (Loureiro et al., 2007). To minimize the artificial variations that may be caused by experimental processes (e.g. Bennett et al., 2008; Doležel and Bartoš, 2005; Nix et al., 2024; Noirot et al., 2005; Praca-Fontes et al., 2011; Trávníček et al., 2015), including methodological and plant environmental variables, five individuals were randomly collected from common gardens at the same growth stage. Further, the internal standard was always from the same tomato plant; all samples were measured within 2 h of removal from the plant; all measurements were conducted by one person (Qing-Song Xiao); only CyStain^TM^ from a single batch was used for staining; all estimations were performed in a CyFlow Space flow cytometer. All estimations cannot be completed with a day. Variations may be caused by different-day conditions, such as room temperature and degree of chopping. To exclude impact of artificial variations, we selected ten individuals of *R. tibetica* to estimates GS three times on three different days. The CV is very low: 0.176%-1.445% (Table S4), indicating the reliability of GS estimation. Based on the result of GS estimation, there is no polyploid in this species.

### 2.3 Leaf trait measurements

About 6–10 individual leaves were collected from each population sampled. A total of 218 individual leaves were collected. The leaves were dried with silica gel, flattened in a small penetrative bag, and transported to the laboratory. In total, 150 leaves were collected from common gardens and were also dried with silica gel. Leaves were trimmed using scissors to measure stomatal size and number. The middle part of the leaf was retained and soaked in double-distilled water for 2–3 days. The soaked leaves were placed on glass slides and the stomata of five different fields of view on the abaxial and adaxial sides of the leaves were observed and photographed using a fluorescence microscope (Leica DM5000B). The photographs were imported into ImageJ (Rueden et al., 2017) to measure the size and number of stomata on the abaxial and adaxial sides of the leaves. As the stomatal number for all samples was measured in the same visual field and scale, it can also serve as a measure of stomatal density.

### 2.4 Environmental data collection

The climate data, including monthly mean precipitation (Prec, mm), solar radiation (Srad, kJ m^-2^ day^-1^), monthly mean temperature (Tavg, °C) as well as water vapor pressure (Vapr, kPa) from January to December, were downloaded from the WorldClim version 2.1 with a resolution of 2.5 minutes (https://www.worldclim.org) (Fick and Hijmans, 2017). Soil data with a resolution of 30 arcseconds were downloaded from the Land-Atmosphere Interaction Research Group at Sun Yat-sen University website (http://globalchange.bnu.edu.cn/research/soil2) (Shangguan et al., 2013). Soil data included alkali-hydrosoluble nitrogen (AN, mg/kg), available phosphorus (AP, mg/kg), available potassium (AK, mg/kg), total nitrogen (TN, g/100 g), total phosphorus (TP, g/100 g), total potassium (TK, g/100 g), and soil organic matter (SOM, g/100 g).

### 2.5 Statistical analyses

The significance of the differences in GS among the three ecotypes was examined using the t-test functions “ggplot” and “geom_signif” in the R packages *ggplot2* (Wickham, 2016) and *ggsignif* (Ahlmann-Eltze and Patil, 2021), respectively, with *p* < 0.05, indicating a significant difference. We performed Pearson’s correlation analyses in the *corrplot* (Simko, 2021) and *ggplot* packages to estimate the relationships between intraspecific GS and geographical factors (altitude, longitude, latitude), climatic factors (Prec, Srad, Tavg, Vapr) and soil nutrients (AN, AP, AK, TN, TP, TK, SOM). Further, *p* < 0.05, *p* < 0.01, *p* < 0.001 indicated the significance levels of Pearson’s correlations. To explore which climate factors can predict intraspecific GS variation in *R. tibetica*, we performed regression subset selection by exhaustive search using the “regsubsets” function in the *leaps* package (Lumley, 2020). This function calculated adjust-R^2^ for different generalized linear models (GLM) based on different climatic factors. The best model was selected using adjust-R^2^, where the higher the value, the better the model. Bayesian information criterion (BIC) was then used to select the optimal and simplest model through an all-subset regression analysis using the *leaps* package. The optimal model was indicated by the lowest BIC, and adjust-R^2^ with *p* value < 0.05 was used to support the model. To validate the predictions of “regsubsets” and determine the relative contribution of climatic factors to GS, a relative weight analysis was conducted using the “relWeights” function (https://rdrr.io/github/jgodet/utilitR/man/relWeights.html). For both stepwise regression and relative weight analysis, GS served as the dependent variable and climatic factors as the independent variables. All analyses were performed using R version 4.0.3 (R Core Team, 2021).

To test the influence of geography and environment on intraspecific GS variation, we performed Mantel and partial Mantel tests with the R package *vegan* (Oksanen et al., 2018) using the “mantel” and “mantel.Partial” functions. To eliminate multicollinearity among factors, we performed principal component analysis (PCA) on altitude and climate factors using the functions “prcomp” and “ggplot” in the *ggplot2* package, and then used the first and second principal component data to calculate the environmental parameter distance between populations. Based on the longitude and latitude information of each population, the geographic Euclidean distance between populations was calculated using the R package *geosphere* (Hijmans, 2019) with the function “distm”. For the partial Mantel tests, geography and environment were controlled separately.

We analyzed Pearson’s correlations between stomatal traits from wild and common gardens and intraspecific GS using the R packages *ggplot2* and *ggpubr* (Kassambara, 2020). The “corrplot” function in the R package *corrplot* was used to test the relationships between morphological traits and environmental factors using Pearson’s correlations. These correlations were judged by the correlation coefficients (*r*) and *p* values, where *p* < 0.05 indicated a significant correlation.

To explore the association between of stomatal traits variation and GS, pair-wised t-test of stomatal traits between wild and common garden populations were estimated using the R function *t.test*. If *t* is a negative (positive) number and *p* < 0.05, indicating stomatal traits of wild populations are significantly less (more) than those traits of common garden populations. Then, the correlations between absolute value of differences of stomatal traits (traits of wild populations minus traits of common garden populations) and GS were estimated by *ggpmisc* and *ggpubr*.

## 3. Results

### 3.1 Estimation of GS

This study measured GS in 53 populations, and included 265 individuals of *R. tibetica* (Tables S1, S2). The mean GS was approximately 1956.76 Mb, ranging from 1892.59 to 2018.32 Mb with about 1.066-fold variation (Fig. 1 c). The GS values from all three ecotypes declined linearly with increasing altitude (Fig. 2 a). The mean GS of the LF ecotype was the largest (1984.00 ± 30.39 Mb), followed by that of HF (1942.40 ± 27.89 Mb), and AM had the smallest value (1937.40 ± 30.50 Mb) (Fig. 2). Moreover, the GS of the LF group was significantly higher than those of the HF and AM groups (*p* < 0.001; Fig. 2 b). There was no significant difference in the GS between the two high-elevation ecotypes, HF and AM.

**Fig. 2.**
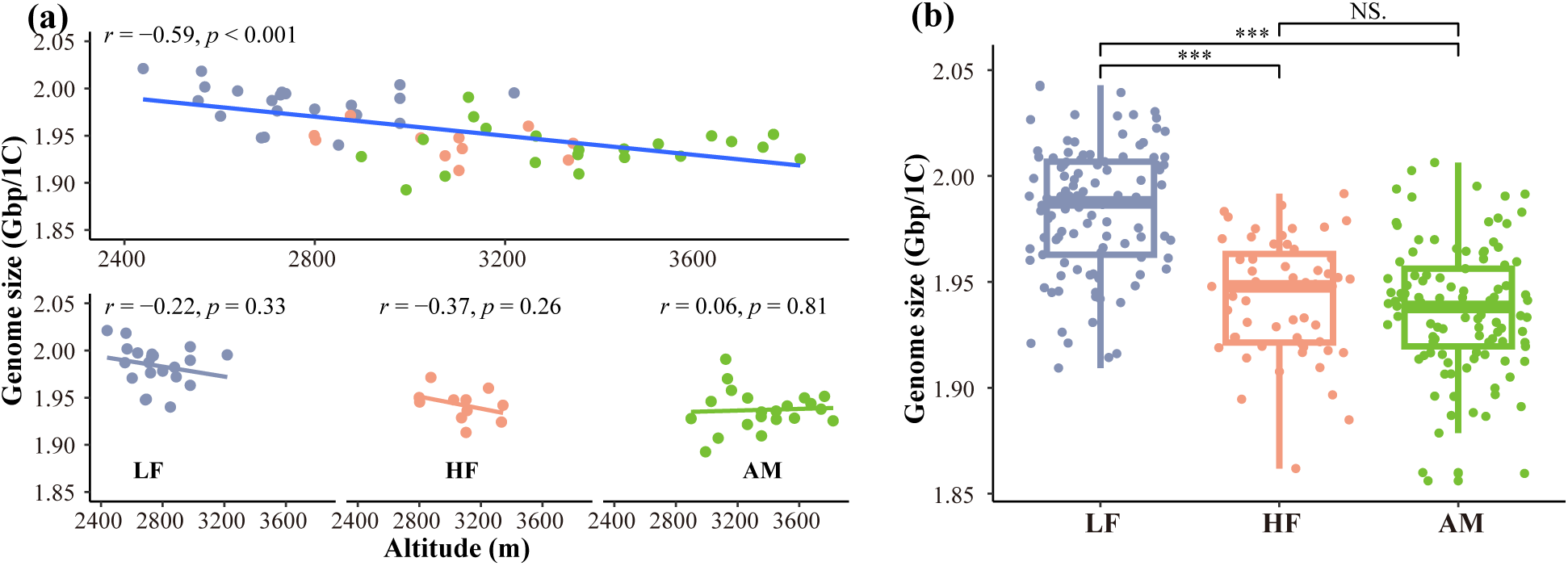
Genome size (GS) variation of wild *R. tibetica* populations with altitude. (a) correlations between entire GS and altitude (above) and between GS within each ecotype and altitude (below). *r* is the correlation coefficient and *p* indicates the significant level of *r*. (b) differences in GS among the three ecotypes. AM, alpine meadow type; HF, high-altitude forest type; LF, low-altitude forest type. Significance levels are denoted by asterisks: \*\*\**p* < 0.001; NS = non-significant.

### 3.2 Correlation between GS and environmental factors

Overall, there was a significant positive correlation between GS and monthly climatic factors (Fig. S1). GS was positively correlated with monthly mean temperature (*r* = 0.66, *p* < 0.001), monthly mean precipitation (*r* = 0.57, *p* < 0.001), and water vapor pressure (*r* = 0.63, *p* < 0.001; Fig. 3 a–d). However, solar radiation in June, July, and August, were negatively correlated with GS and only in July, the correlation was significant (*r* = -0.42, *p* < 0.01; Fig. S2). Since the growing season for *R. tibetica* consists of June, July, and August, the growing season solar radiation included that of only these three months for the following analysis. The results indicated that GS was significantly negatively correlated with solar radiation during the growing season (*r* = -0.32, *p* < 0.05; Fig. 3 a and e). Our results showed a significant negative correlation between GS and altitude (*r* = -0.59, *p* < 0.001; Fig. 2 a). within ecotypes, the correlations between GS and altitude were not significant. In addition, a significant negative correlation was found between the GS of *R. tibetica* and latitude (*r* = -0.49, *p* < 0.001; Fig. 3 a). There was no correlation between GS and longitude or soil nutrients (Fig. 3 a). In addition, with increasing altitude, precipitation, temperature, and water vapor pressure decreased, while solar radiation and soil nutrients increased (Fig. 3 a). For stomatal traits from the wild and common garden conditions, no significant correlations were found between stomatal size and density on either the abaxial or adaxial sides of the leaves and environmental factors except significant correlations between stomatal size and precipitation (*p* < 0.01; Fig. S3).

**Fig. 3.**
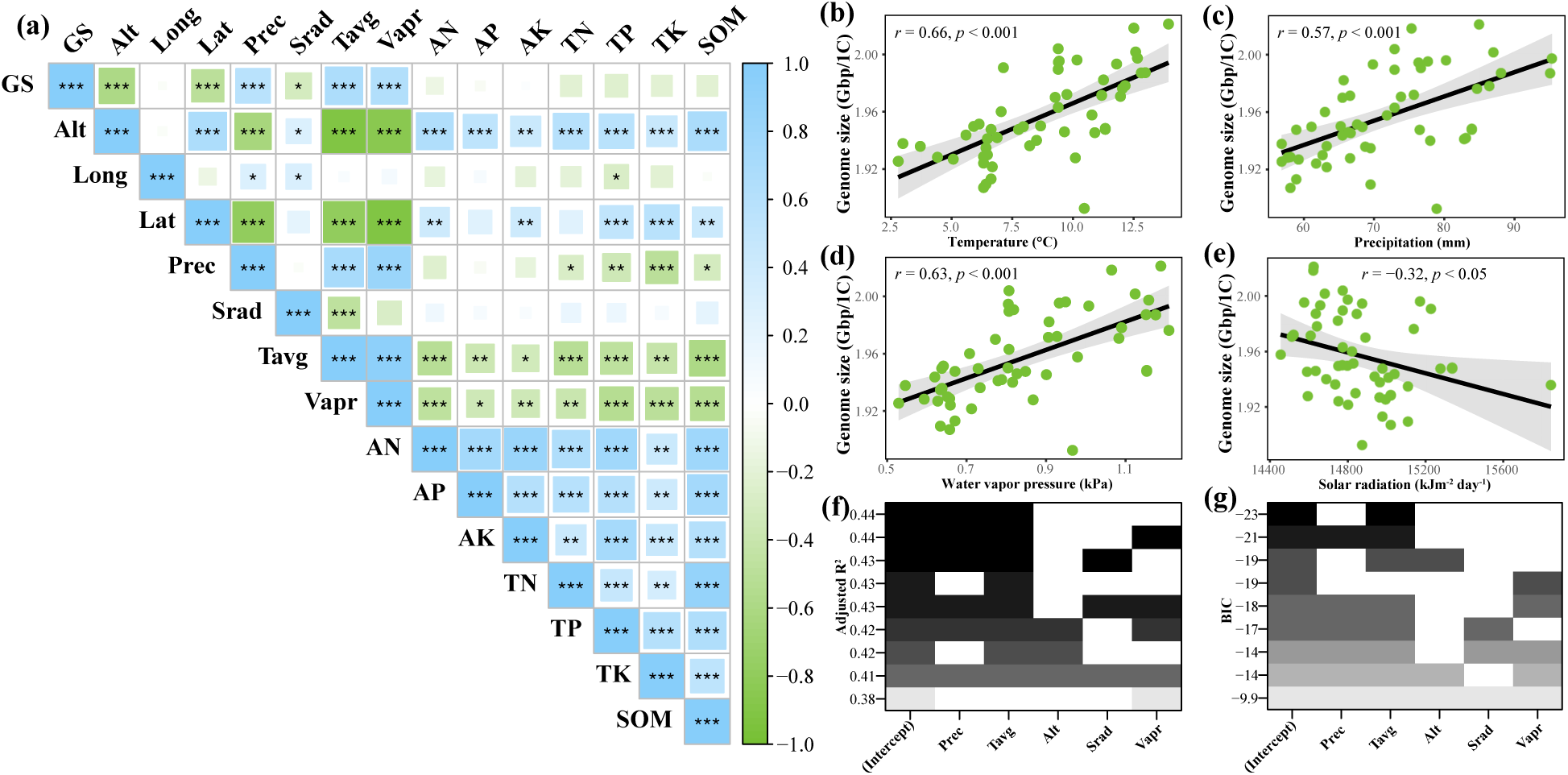
Correlations between genome size (GS) and environmental factors for *R. tibetica*. (a) Pearson’s correlation coefficients among GS and geographic factors (altitude, longitude, latitude), climatic factors (Prec, Srad, Tavg, Vapr), and soil nutrients (AN, AP, AK, TN, TP, TK, SOM). Srad is the solar radiation of the growing season from June to August. Significance levels are denoted by asterisks: \**p* < 0.05; \*\**p* < 0.01; \*\*\**p* < 0.001. Alt, altitude; Long, longitude; Lat, latitude; Prec, monthly mean precipitation; Srad, solar radiation of growth season from June to August; Tavg, monthly mean temperature; Vapr, water vapor pressure; AN, alkali-hydrosoluble nitrogen; AP, available phosphorus; AK, available potassium; TN, total nitrogen; TP, total phosphorus; TK, total potassium; SOM, soil organic matter. (b-e) scatter plots showing the relationship between GS and four climatic factors. The black regression lines represent the least-squares estimates of the conditional mean function, and the 95% confidence interval is shown by the grey shading around the line. Model selection through Adjust-R^2^ estimated by regression (f) and Schwarz’s information criterion (BIC) (g).

Regression subset analysis indicated that precipitation + temperature and precipitation + temperature + water vapor pressure were two equally good models (both had adjusted R^2^ = 0.44; Fig. 3 f) that best explained the GS variation. The lowest BIC value, further indicated that temperature was the best independent variable to explain GS variability (multiple R^2^ = 0.44, adjusted R^2^ = 0.43, df = 51, *p* < 0.001; Fig. 3 g). Relative weight analysis suggested that precipitation (R^2^ = 0.25) and temperature (R^2^ = 0.24) contributed relatively more to GS variation, followed by water vapor pressure (R^2^ = 0.21). The total R^2^ value for all the independent variables together was 0.47. These analyses suggest that precipitation and temperature can explain the GS variation well.

### 3.3 Contribution of geography and environmental factors to GS variation

Mantel tests showed a strong positive correlation between GS and both geographic distance (Mantel statistic *r* = 0.178, *p* < 0.01) and environmental factors (*r* = 0.290, *p* < 0.001) (Table 1). In the partial Mantel tests, when the environment was controlled for, GS exhibited no significant association with geographic distance (*r* = -0.004, *p* > 0.05), whereas it remained positively correlated with the environmental factors when geographic distance was controlled for (*r* = 0.233, *p* < 0.001) (Table 1).

**Table 1.** The relationships between genome size variation and geography or environment.

| Models | Geography |  | Environments |  |
| --- | --- | --- | --- | --- |
|  | *r* | *p* | *r* | *p* |
| **Mantel tests** |  |  |  |  |
| Genome size | **0.178** | **< 0.01** | **0.290** | **< 0.001** |
| **Partial Mantel tests** |  |  |  |  |
| Environments were controlled | -0.004 | 0.513 | - |  |
| Geography was controlled | - |  | **0.233** | **< 0.001** |

### 3.4 Analysis of the relationship between morphological traits and GS

For *R. tibetica* collected from the wild, stomatal size on both the abaxial and adaxial sides of the leaves showed positive correlations with GS (*r* = 0.52, *p* < 0.05, and *r* = 0.52, *p* < 0.05, respectively) (Fig. 4 a). Stomatal density at the abaxial of the leaves was negatively correlated with GS (*r* = -0.47, *p* < 0.05), while the correlation between stomatal density at the adaxial of the leaves and GS was not significant (*r* = -0.32, *p* > 0.05) (Fig. 4 b). For plants cultivated in the common garden, no significant correlations between stomatal size and density on either the abaxial or adaxial sides of the leaves and GS were detected (Fig. 4 a and b).

**Fig. 4.**
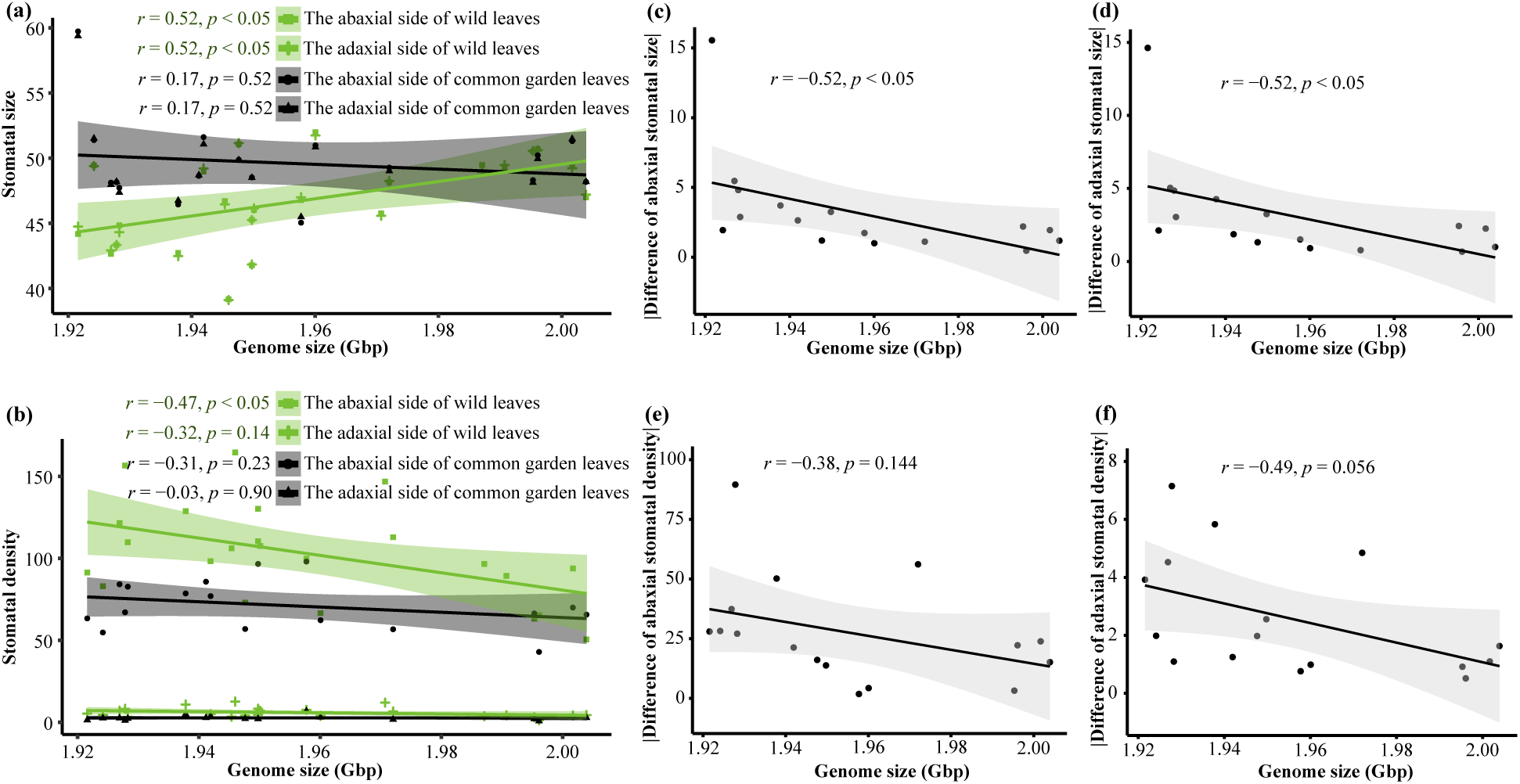
The relationships between genome size (GS) and stomatal traits. (a) and (b) are correlations between GS and stomatal size, and stomatal density respectively. Green represents the traits sampled from the wild and gray represents the traits sampled from common gardens. The regression lines represent the least-squares estimate of the conditional mean function, and 95% confidence interval is shown by the shading around the line. Stomatal size and stomatal density of the abaxial and adaxial surfaces of leaves were measured. (c-d) correlations between GS and absolute stomatal differences (traits of wild populations minus traits common garden populations).

It was important that stomatal size from wild populations was significantly less than that from common garden populations (abaxial side: *t* = −2.25, *p* < 0.05; adaxial side: *t* = −2.24, *p* < 0.05). The stomatal density from wild populations was significantly more than that from common garden populations (abaxial side: *t* = 3.98, *p* < 0.01; adaxial side: *t* = 5.01, *p* < 0.001). More importantly, the absolute values of differences of stomatal size for both sides were negatively and significantly correlated with GS (both *r* = −0.52, *p* < 0.05; Fig. 4 c and d). Although the correlations between absolute differences of stomatal density and GS were not significant, the relationships were negative (abaxial side: *r* = −0.38, *p* = 0.144; adaxial side: *r* = −0.49, *p* = 0.056; Fig. 4 e and f).

## 4. Discussion

### 4.1 Smaller genome size promotes Roscoea tibetica to adapt to higher elevation

Although the effect of environmental stress on GS evolution has been extensively studied, the role of environmental factors in intraspecific GS variation remains poorly understood (Blommaert, 2020; Faizullah et al., 2021). A recent study revealed that the distribution of angiosperm GS is shaped by climate (Bureš et al., 2024), suggesting that climatic factors play a crucial role in the evolution of GS. Our analysis suggests that variations in intraspecific GS in *R. tibetica* across the Hengduan Mountains were determined by environmental factors rather than randomly distributed. Previous research suggests that environmental pressures are the direct driving forces of natural selection and evolutionary change (Hoffmann and Hercus, 2000). If the evolution of the GS is neutral in *R. tibetica*, there is no significant association between the GS and environmental factors. The intraspecific GS of *R. tibetica* was significantly related to altitude, latitude, monthly precipitation, monthly average temperature, water vapor, and solar radiation (Fig. 2 and 3). The Mantel test indicates that the environment shapes the geographical pattern of GS variation in *R. tibetica* (Table 1). Among the environmental factors, precipitation and temperature were the best predictors of GS variation, suggesting their crucial role in the evolution of intraspecific GS. Thus, the close relationships between intraspecific GS and environmental factors suggest that the variations observed across the distribution range of *R. tibetica* are the result of adaptive evolution. This observation mirrors a similar pattern observed in maize, which likely reflects the effect of natural selection on GS variation (Bilinski et al., 2018; Diez et al., 2013).

Environmentally stressed conditions, such as low CO_2_, water, and nutrient levels, may favor species with smaller genomes (Faizullah et al., 2021). Plants with small genomes are more adaptable to harsh environments (Leitch and Leitch, 2013), can maximize water-use efficiency and CO_2_ absorption, and facilitate rapid growth in dry or variable water environments (Hetherington and Woodward, 2003; Simonin and Roddy, 2018; Záveská et al., 2024). We found that for *R. tibetica*, the GS significantly decreased with increasing altitude and solar radiation during the growing season and increased with increasing precipitation, temperature, and water vapor pressure (Fig. 2 and 3). The stomatal size of *R. tibetica* shows a positive correlation with GS for the wild populations. The closing and opening of stomata are slower for individuals with larger stomatal sizes (Kardiman and Raebild, 2018; Lawson and Matthews, 2020), which may hinder efficient water management in drier environments (Šmarda et al., 2023; Veselý et al., 2020). This speculation is consistent with the large genome constraint hypothesis (Bureš et al., 2024; Knight et al., 2005). Higher altitude, lower precipitation, temperature, and water vapor, and stronger solar radiation likely created environmentally stressed conditions that influenced the evolution of GS in *R. tibetica*. These conditions enabled plants with small GS to survive in the harsher environment along the elevation.

More importantly, the results supported our assumption based on the large genome constraint hypothesis that the smaller of the GS exhibit larger variation of stomatal traits between populations from the wild and common garden (Fig. 4 c-f). The large genome constraint hypothesis emphasized the effects of GS on minimum cell size (Bennett, 1987; Bhadra et al., 2023; Šmarda et al., 2023; Theroux-Rancourt et al., 2021; Veselý et al., 2020). This hypothesis gives species or individuals with smaller GS have greater flexibility in GS-dependent traits, compared to those with larger genomes. Stomatal size and density are two main traits correlated with GS variation (Beaulieu et al., 2008; Faizullah et al., 2021; Lomax et al., 2014; Roddy et al., 2020; Simonin and Roddy, 2018; Veselý et al., 2020). Common garden in our study provided a more comfortable living condition for individuals of *R. tibetica* compared with the wild conditions especially for higher-elevational individuals. *Roscoea tibetica* has larger stomatal size and lower stomatal density in the common garden than those traits in the wild (Fig. 4). Under such situation, degree of variation of the two traits between the wild and common garden populations are negatively correlated with GS (Fig. 4 c-f), indicating populations of *R. tibetica* with smaller GS from higher elevation exhibit larger variations of stomatal size and density. This result suggested populations with smaller GS are more flexible to respond to habitat change through large-scale changes in morphological traits, which is likely an adaptive feature for *R. tibetica* to thrive in higher elevation.

Moreover, several studies have proposed that the genome streamlining hypothesis holds true in environments with extremely limited amounts of P and N (Bales and Hersch-Green, 2019; Faizullah et al., 2021; Kang et al., 2015). However, our study found no correlation between GS and soil nutrients, which does not support the genome streamlining hypothesis. For *R. tibetica*, soil nutrients in the habitats increased with increasing altitude (Fig. 3 a). This finding suggests that the soil nutrients in the habitats were sufficient and did not create stressful conditions, thereby exerting a weak or no effect on GS changes in *R. tibetica*. A potential causal explanation could be that rhizomes, functioning as storage organs in Zingiberaceae, prevent plants from being affected by nutrient deficiency (Záveská et al., 2024).

### 4.2 Stomatal traits do not reliably predict intraspecific GS variations

Many studies indicate that GS is significantly positively correlated with stomatal size and negatively correlated with stomatal density (Beaulieu et al., 2008; Faizullah et al., 2021; Lomax et al., 2014; Roddy et al., 2020; Simonin and Roddy, 2018; Veselý et al., 2020). Current models suggest that selection for the GS is influenced by stomatal size (Faizullah et al., 2021), implying that environmental selection based on stomatal size may influence GS evolution in plants (Veselý et al., 2020). However, our findings challenge this traditional understanding, as our observations in the wild clearly indicate that GS is positively correlated with stomatal size and negatively correlated with stomatal density (Fig. 4). However, these significant correlations disappeared when plants from the common garden were analyzed, strongly suggesting the plasticity of stomatal traits. The plasticity of stomatal traits has been reported in other plants (Sun et al., 2014; Wang et al., 2019), suggesting that stomatal traits are controlled by environmental factors rather than the GS. A growth chamber experiment of highland teosintes revealed weak support for a positive correlation between GS and cell size (Bilinski et al., 2018), suggesting that the size relationship could be impacted by habitats. Comprehensive evolutionary analysis of the GS and stomatal size in Proteaceae revealed that ancient changes in the GS clearly impacted stomatal size in Proteaceae, but adaptive responses to habitat strongly changed the genome–stomatal size relationship, suggesting that environmental adaptation in stomatal size is independent of the effects of the GS (Jordan et al., 2015). Thus, Jordan et al. (2015) proposed that a general genome–stomatal size relationship cannot be expected through geological time because habitats and other factors, such as atmospheric CO_2_ concentration, could influence stomatal size independent of GS. Previous studies, along with our findings on stomatal trait differences between wild and common gardens, strongly suggest that stomata may not reliably predict intraspecific GS variations, especially for the transplanted species.

Both the GS and stomatal size are affected by environmental factors. Thus, the formation of a causal relationship between stomatal traits in the wild and the GS was likely dominated by environmental factors. Moreover, the decoupling of the genome–stomatal size relationship observed when analyzing traits from common gardens suggests that selection pressures are likely reduced in controlled environments, such as common gardens, providing the opportunity for the smaller-genome populations to relax the limitation of stomatal traits to adapt to new habitat.

## 5. Conclusions

This study sheds light on the adaptive evolution of intraspecific GS variation in the alpine herb *R. tibetica* from two perspectives. First, environmental factors, especially precipitation and temperature, are the main stressors that cause plants with large genome sizes to survive in harsh environments at high altitudes. Second, stomatal traits do not reliably predict intraspecific GS variations observed in wild and common garden experiments, suggesting that the nucleotypic and nucleoskeletal hypotheses may not be fully supported by genome–stomatal size relationship. In general, these results for the first time to illustrate the large-genome constraint hypothesis is applicable in the adaptive evolution of intraspecific GS and small GS is beneficial to alpine plant to live in the hasher environments. Although our inferences were just from one species, our findings updated our understandings on the evolution of intraspecific GS. These findings also suggest that *R. tibetica* serves as a valuable model for further exploration of genomic signals for intraspecific GS variation and how GS promotes adaptive evolution in a heterogeneous mountainous environment.

## Data accessibility statement

All data for this work are available in Table S1-S4.

## CRediT authorship contribution statement

**Jian-Li Zhao** designed the study. **Jian-Li Zhao**, **Li Li**, **Qing-Song Xiao** and **Hong-Fan Chen** collected samples from the wild. **Qing-Song Xiao** collected the genome size data. **Wen Guo** contributed genome size data collection using flow cytometric measurement. **Hong-Fan Chen** collected the data of stomatal traits. **Qing-Song Xiao** performed the analyses. **Jian-Li Zhao** and **Li Li** contributed to analyses. **Qing-Song Xiao** and **Jian-Li Zhao** drafted the first version of the manuscript. **Tomáš Fér** deeply improved the draft from data analysis and writing. **Qing-Song Xiao**, **Tomáš Fér**, **Wen Guo**, **Li Li** and **Jian-Li Zhao** contributed to interpretation of the results, and the final form of the manuscript; All authors gave final approval for publication and agreed to be held accountable for the work performed therein.

## Acknowledgments

We thank Wen-Jing Wang and Guo-Jun Shao from Yunnan University for their help during our sampling. This work was supported by the National Natural Science Foundation of China (grant number 41871047, 32101355), the “Young Talent Project” of Yunnan (grant number YNWR-QNBJ-2019-214), the Project for Talent and Platform of Science and Technology in Yunnan Province Science and Technology Department (grant number 202205AM070005).

