## Supplementary material for "The small genome size ensures adaptive flexibility for an alpine ginger": Table S1

| **Table S1** Sampling information and average genome size of each population for *Roscoea tibetica*. | | | | | | |
| --- | --- | --- | --- | --- | --- | --- |
| Populations | Longitude | Latitude | Altitude (m) | GS (Mb) | SD | Ecotypes |
| RT1 | 99.648185 | 24.174006 | 2555 | 1987.08 | 24.75 | LF |
| RT2 | 99.643036 | 24.176433 | 2638 | 1997.38 | 21.78 | LF |
| RT3 | 99.3020845 | 25.1663595 | 2439.5 | 2021.13 | 15.28 | LF |
| RT4 | 99.105855 | 25.849149 | 2710 | 1987.24 | 19.48 | LF |
| RT5 | 99.104521 | 25.866695 | 2800 | 1978.15 | 22.54 | LF |
| RT6 | 99.105855 | 25.889055 | 2569 | 2001.68 | 18.65 | LF |
| RT7 | 99.320269 | 26.456769 | 3219 | 1995.32 | 18.17 | LF |
| RT8 | 99.440519 | 26.583715 | 2888 | 1972.03 | 27.89 | LF |
| RT9 | 99.594498 | 26.573175 | 2562 | 2018.32 | 17.05 | LF |
| RT31 | 102.349119 | 27.646448 | 2850 | 1940.05 | 35.39 | LF |
| RT32 | 102.90724 | 24.758075 | 2721 | 1976.25 | 27.87 | LF |
| RT33 | 102.928076 | 24.762693 | 2688 | 1947.71 | 25.54 | LF |
| RT34 | 102.930114 | 24.763735 | 2694 | 1948.36 | 24.60 | LF |
| RT38 | 101.738956 | 27.542409 | 2979 | 2003.92 | 16.67 | LF |
| RT39 | 101.738956 | 27.542409 | 2979 | 1963.04 | 20.41 | LF |
| RT40 | 101.738956 | 27.542409 | 2979 | 1989.57 | 16.86 | LF |
| RT41 | 101.978531 | 28.339439 | 2739 | 1994.54 | 34.95 | LF |
| RT45 | 100.789154 | 26.769254 | 2877 | 1982.21 | 31.81 | LF |
| RT48 | 100.143299 | 25.645859 | 2602 | 1970.71 | 30.39 | LF |
| RT49 | 101.229057 | 25.951408 | 2728 | 1993.27 | 17.57 | LF |
| RT54 | 103.371015 | 26.099017 | 2731 | 1996.10 | 25.56 | LF |
| RT12 | 100.211535 | 27.017996 | 2802 | 1945.39 | 16.42 | HF |
| RT13 | 100.228465 | 27.031988 | 2875 | 1971.42 | 16.78 | HF |
| RT14 | 99.9693845 | 27.3609245 | 2799 | 1950.10 | 30.40 | HF |
| RT16 | 99.805984 | 27.566113 | 3249 | 1960.07 | 23.51 | HF |
| RT21 | 100.023823 | 27.687784 | 3333 | 1924.16 | 21.50 | HF |
| RT23 | 100.026226 | 27.557591 | 3110 | 1936.27 | 24.24 | HF |
| RT27 | 99.753113 | 28.141027 | 3074 | 1928.67 | 43.22 | HF |
| RT28 | 99.773884 | 28.072435 | 3103 | 1947.69 | 26.49 | HF |
| RT29 | 99.773884 | 28.072435 | 3103 | 1913.11 | 23.42 | HF |
| RT50 | 103.243896 | 26.276609 | 3343 | 1941.93 | 26.48 | HF |
| RT53 | 103.35457 | 27.411283 | 3023 | 1947.63 | 15.25 | HF |
| RT10 | 100.18261 | 26.949311 | 3028 | 1945.99 | 18.75 | AM |
| RT11 | 100.2069 | 27.02136 | 2898 | 1927.82 | 23.73 | AM |
| RT15 | 99.8138595 | 27.460168 | 3263.5 | 1921.58 | 34.09 | AM |
| RT18 | 99.615355 | 27.92837 | 3569 | 1928.27 | 30.71 | AM |
| RT19 | 99.910526 | 27.795313 | 3451 | 1926.91 | 24.14 | AM |
| RT20 | 99.983826 | 27.72198 | 3634 | 1949.76 | 20.78 | AM |
| RT22 | 100.623651 | 27.686264 | 3353 | 1929.94 | 16.81 | AM |
| RT24 | 99.987173 | 28.236725 | 3820 | 1925.46 | 19.55 | AM |
| RT25 | 99.866238 | 28.178636 | 3742 | 1937.81 | 24.14 | AM |
| RT26 | 99.753113 | 28.141027 | 3074 | 1907.11 | 43.39 | AM |
| RT35 | 101.161423 | 28.122785 | 3764 | 1951.37 | 10.73 | AM |
| RT36 | 101.16344 | 28.13855 | 3676 | 1943.72 | 27.84 | AM |
| RT37 | 101.22372 | 27.68637 | 3265 | 1949.60 | 23.44 | AM |
| RT42 | 101.398144 | 29.092477 | 3355 | 1934.83 | 19.98 | AM |
| RT43 | 101.398144 | 29.092477 | 3355 | 1909.49 | 17.40 | AM |
| RT44 | 100.995941 | 27.330295 | 3134 | 1970.03 | 26.21 | AM |
| RT47 | 100.361824 | 25.971472 | 3160 | 1957.73 | 23.78 | AM |
| RT51 | 103.248207 | 26.248128 | 3522 | 1941.17 | 26.57 | AM |
| RT52 | 103.322651 | 27.406139 | 3123 | 1990.73 | 17.94 | AM |
| RT55 | 102.826598 | 26.039543 | 2992 | 1892.59 | 24.47 | AM |
| RT56 | 97.564751 | 28.794293 | 3450 | 1935.85 | 16.25 | AM |
| GS, population-average genome size estimates of 5 individuals; SD, standard deviation. | | | | | | |
| LF, low-altitude forest type; HF, high-altitude forest type; AM, alpine meadow type. | | | | | | |
