## Supplementary material for "The small genome size ensures adaptive flexibility for an alpine ginger": Table S2

| **Populations** | **Numbers** | **Sample numbers** | **Mean-X** | | **CV-X%** | | **Count** | | **C-value/Mb** |
| --- | --- | --- | --- | --- | --- | --- | --- | --- | --- |
|  |  |  | **Tomato** | ***R. tibetica*** | **Tomato** | ***R. tibetica*** | **Tomato** | ***R. tibetica*** |  |
| RT1 | 1 | RT1-1 | 72.09 | 149.33 | 3.55 | 3.00 | 6762 | 6615 | 1984.438 |
|  | 2 | RT1-2 | 76.08 | 155.67 | 3.82 | 3.75 | 15052 | 8606 | 1960.198 |
|  | 3 | RT1-3 | 78.18 | 164.41 | 3.67 | 2.96 | 3631 | 5093 | 2014.643 |
|  | 4 | RT1-4 | 79.54 | 166.88 | 4.33 | 3.35 | 5263 | 4499 | 2009.945 |
|  | 5 | RT1-5 | 86.34 | 177.20 | 3.07 | 3.06 | 4261 | 6952 | 1966.152 |
| RT2 | 1 | RT2-1 | 76.53 | 159.01 | 3.47 | 2.83 | 11778 | 5481 | 1990.482 |
|  | 2 | RT2-2 | 81.55 | 167.67 | 2.95 | 2.68 | 3970 | 5950 | 1969.686 |
|  | 3 | RT2-3 | 87.64 | 183.79 | 3.76 | 2.80 | 3427 | 6550 | 2009.024 |
|  | 4 | RT2-4 | 76.23 | 161.32 | 3.97 | 3.05 | 6748 | 5859 | 2027.346 |
|  | 5 | RT2-5 | 73.28 | 152.25 | 3.73 | 3.69 | 8923 | 7205 | 1990.386 |
| RT3 | 1 | RT3-1 | 82.89 | 175.35 | 4.32 | 3.64 | 3534 | 3839 | 2026.605 |
|  | 2 | RT3-2 | 61.14 | 129.45 | 4.39 | 3.74 | 8803 | 5821 | 2028.346 |
|  | 3 | RT3-3 | 69.57 | 148.10 | 4.10 | 3.84 | 3003 | 5911 | 2039.382 |
|  | 4 | RT3-4 | 69.61 | 145.41 | 3.10 | 2.83 | 10181 | 11520 | 2001.189 |
|  | 5 | RT3-5 | 74.81 | 156.97 | 3.42 | 2.94 | 6556 | 7309 | 2010.122 |
| RT4 | 1 | RT4-1 | 66.66 | 139.53 | 3.72 | 3.36 | 4361 | 6408 | 2005.247 |
|  | 2 | RT4-2 | 75.28 | 155.46 | 3.90 | 3.40 | 8569 | 6376 | 1978.357 |
|  | 3 | RT4-3 | 80.49 | 168.97 | 3.68 | 3.46 | 7877 | 6720 | 2011.098 |
|  | 4 | RT4-4 | 54.65 | 112.47 | 4.92 | 4.73 | 6443 | 5104 | 1971.569 |
|  | 5 | RT4-5 | 71.08 | 146.16 | 3.96 | 3.76 | 10489 | 6419 | 1969.911 |
| RT5 | 1 | RT5-1 | 95.69 | 195.10 | 3.33 | 3.07 | 7526 | 4310 | 1953.243 |
|  | 2 | RT5-2 | 62.52 | 130.04 | 4.31 | 3.81 | 9645 | 4357 | 1992.615 |
|  | 3 | RT5-3 | 69.82 | 142.39 | 4.36 | 3.48 | 17035 | 9430 | 1953.733 |
|  | 4 | RT5-4 | 74.13 | 154.40 | 3.26 | 2.83 | 8445 | 6079 | 1995.349 |
|  | 5 | RT5-5 | 66.99 | 139.56 | 4.27 | 3.68 | 9751 | 6679 | 1995.798 |
| RT6 | 1 | RT6-1 | 78.85 | 163.26 | 3.48 | 3.38 | 7768 | 5496 | 1983.552 |
|  | 2 | RT6-2 | 77.49 | 162.20 | 3.74 | 3.18 | 5997 | 6630 | 2005.260 |
|  | 3 | RT6-3 | 84.61 | 178.63 | 4.16 | 3.17 | 8456 | 6360 | 2022.545 |
|  | 4 | RT6-4 | 86.15 | 178.17 | 2.93 | 2.69 | 3711 | 5758 | 1981.275 |
|  | 5 | RT6-5 | 78.74 | 165.68 | 3.54 | 3.04 | 5770 | 10685 | 2015.766 |
| RT7 | 1 | RT7-1 | 82.72 | 172.93 | 4.02 | 3.61 | 4581 | 2466 | 2002.743 |
|  | 2 | RT7-2 | 73.39 | 154.73 | 4.41 | 3.53 | 5279 | 6858 | 2019.776 |
|  | 3 | RT7-3 | 80.53 | 165.70 | 3.48 | 2.78 | 7586 | 5665 | 1971.198 |
|  | 4 | RT7-4 | 77.88 | 162.32 | 3.16 | 2.82 | 5780 | 5557 | 1996.694 |
|  | 5 | RT7-5 | 72.21 | 149.71 | 3.39 | 3.05 | 10249 | 8080 | 1986.182 |
| RT8 | 1 | RT8-1 | 76.68 | 155.86 | 4.62 | 3.51 | 9187 | 5876 | 1947.234 |
|  | 2 | RT8-2 | 72.72 | 151.02 | 3.33 | 2.82 | 7132 | 5443 | 1989.510 |
|  | 3 | RT8-3 | 81.56 | 165.42 | 3.99 | 3.05 | 7196 | 6079 | 1943.016 |
|  | 4 | RT8-4 | 78.59 | 164.79 | 3.45 | 3.50 | 10411 | 5114 | 2008.765 |
|  | 5 | RT8-5 | 84.40 | 173.70 | 3.95 | 3.16 | 8548 | 7844 | 1971.618 |
| RT9 | 1 | RT9-1 | 70.92 | 150.20 | 3.68 | 3.61 | 10887 | 5096 | 2028.928 |
|  | 2 | RT9-2 | 76.02 | 162.10 | 3.51 | 3.02 | 3627 | 5091 | 2042.776 |
|  | 3 | RT9-3 | 80.05 | 167.55 | 3.28 | 3.16 | 6056 | 5690 | 2005.158 |
|  | 4 | RT9-4 | 73.64 | 153.96 | 2.97 | 3.63 | 10926 | 5120 | 2002.902 |
|  | 5 | RT9-5 | 80.56 | 169.18 | 3.52 | 3.57 | 10978 | 8548 | 2011.848 |
| RT10 | 1 | RT10-1 | 72.65 | 146.61 | 3.71 | 2.69 | 5002 | 9873 | 1933.274 |
|  | 2 | RT10-2 | 77.25 | 159.53 | 3.62 | 2.92 | 3097 | 5646 | 1978.379 |
|  | 3 | RT10-3 | 78.62 | 158.75 | 3.66 | 3.45 | 11398 | 7525 | 1934.400 |
|  | 4 | RT10-4 | 78.74 | 159.92 | 3.79 | 3.53 | 3433 | 6469 | 1945.687 |
|  | 5 | RT10-5 | 78.97 | 159.77 | 3.70 | 3.26 | 11981 | 5248 | 1938.200 |
| RT11 | 1 | RT11-1 | 76.46 | 154.86 | 3.44 | 3.50 | 7946 | 5363 | 1940.307 |
|  | 2 | RT11-2 | 75.39 | 149.20 | 3.86 | 3.11 | 9177 | 10544 | 1895.923 |
|  | 3 | RT11-3 | 78.80 | 158.08 | 3.55 | 3.01 | 7712 | 5947 | 1921.836 |
|  | 4 | RT11-4 | 77.95 | 159.44 | 3.52 | 2.86 | 4800 | 5471 | 1959.506 |
|  | 5 | RT11-5 | 81.41 | 163.29 | 3.96 | 3.01 | 7947 | 6038 | 1921.531 |
| RT12 | 1 | RT12-1 | 72.23 | 147.57 | 3.77 | 3.45 | 4744 | 9130 | 1957.249 |
|  | 2 | RT12-2 | 77.24 | 157.33 | 3.81 | 2.68 | 5571 | 7130 | 1951.348 |
|  | 3 | RT12-3 | 83.59 | 170.48 | 3.80 | 2.80 | 6228 | 5748 | 1953.820 |
|  | 4 | RT12-4 | 78.82 | 160.26 | 3.87 | 3.09 | 4663 | 6443 | 1947.844 |
|  | 5 | RT12-5 | 71.49 | 143.03 | 3.47 | 3.24 | 9872 | 8466 | 1916.670 |
| RT13 | 1 | RT13-1 | 91.93 | 188.15 | 3.17 | 2.61 | 3123 | 3952 | 1960.706 |
|  | 2 | RT13-2 | 66.75 | 137.62 | 4.24 | 3.58 | 10603 | 6704 | 1975.130 |
|  | 3 | RT13-3 | 74.86 | 155.63 | 3.89 | 3.29 | 9403 | 6595 | 1991.632 |
|  | 4 | RT13-4 | 69.06 | 142.78 | 3.79 | 3.24 | 4744 | 6504 | 1980.643 |
|  | 5 | RT13-5 | 71.20 | 144.85 | 3.92 | 3.46 | 10937 | 8835 | 1948.965 |
| RT14 | 1 | RT14-1 | 62.86 | 127.93 | 4.50 | 3.87 | 7208 | 6623 | 1949.681 |
|  | 2 | RT14-2 | 74.00 | 152.20 | 3.70 | 3.42 | 6842 | 5897 | 1970.373 |
|  | 3 | RT14-3 | 72.35 | 150.00 | 4.58 | 2.97 | 5422 | 6624 | 1986.178 |
|  | 4 | RT14-4 | 73.28 | 148.14 | 3.49 | 3.84 | 11871 | 6407 | 1936.656 |
|  | 5 | RT14-5 | 67.54 | 134.49 | 3.89 | 3.98 | 7941 | 4599 | 1907.631 |
| RT15 | 1 | RT15-1 | 77.01 | 151.65 | 4.27 | 3.34 | 6204 | 6172 | 1886.517 |
|  | 2 | RT15-2 | 83.32 | 167.18 | 3.64 | 2.82 | 5739 | 6456 | 1922.209 |
|  | 3 | RT15-3 | 79.54 | 162.15 | 4.05 | 2.80 | 4668 | 6223 | 1952.976 |
|  | 4 | RT15-4 | 80.17 | 163.85 | 3.65 | 3.10 | 5099 | 7070 | 1957.943 |
|  | 5 | RT15-5 | 84.64 | 166.83 | 3.88 | 3.25 | 12975 | 6563 | 1888.270 |
| RT16 | 1 | RT16-1 | 74.59 | 153.48 | 3.29 | 2.90 | 10100 | 5635 | 1971.227 |
|  | 2 | RT16-2 | 74.21 | 152.75 | 3.80 | 2.67 | 6881 | 5654 | 1971.897 |
|  | 3 | RT16-3 | 73.51 | 152.18 | 3.36 | 3.18 | 7162 | 5718 | 1983.246 |
|  | 4 | RT16-4 | 78.98 | 160.77 | 3.95 | 3.63 | 6114 | 6393 | 1950.084 |
|  | 5 | RT16-5 | 82.67 | 166.02 | 3.74 | 3.22 | 10141 | 6864 | 1923.880 |
| RT18 | 1 | RT18-1 | 72.87 | 149.84 | 3.49 | 3.09 | 5858 | 6335 | 1969.901 |
|  | 2 | RT18-2 | 76.28 | 155.39 | 2.70 | 2.42 | 3869 | 4490 | 1951.542 |
|  | 3 | RT18-3 | 81.20 | 161.03 | 3.88 | 3.25 | 9249 | 6158 | 1899.837 |
|  | 4 | RT18-4 | 85.41 | 170.60 | 4.04 | 3.75 | 4462 | 2329 | 1913.532 |
|  | 5 | RT18-5 | 78.08 | 155.39 | 3.87 | 3.25 | 10834 | 5091 | 1906.553 |
| RT19 | 1 | RT19-1 | 78.84 | 157.97 | 3.78 | 3.11 | 7554 | 7645 | 1919.524 |
|  | 2 | RT19-2 | 79.93 | 162.96 | 3.67 | 3.54 | 7686 | 3386 | 1953.155 |
|  | 3 | RT19-3 | 76.19 | 154.64 | 4.27 | 3.17 | 5308 | 7772 | 1944.417 |
|  | 4 | RT19-4 | 91.63 | 180.88 | 2.97 | 2.59 | 7137 | 2631 | 1891.117 |
|  | 5 | RT19-5 | 72.25 | 145.28 | 3.59 | 3.13 | 5860 | 6018 | 1926.342 |
| RT20 | 1 | RT20-1 | 77.82 | 157.66 | 3.25 | 3.35 | 8834 | 7182 | 1940.867 |
|  | 2 | RT20-2 | 78.28 | 161.52 | 4.47 | 3.01 | 4049 | 5762 | 1976.701 |
|  | 3 | RT20-3 | 83.43 | 167.57 | 3.20 | 2.91 | 2360 | 5956 | 1924.153 |
|  | 4 | RT20-4 | 75.54 | 153.18 | 3.63 | 3.31 | 9683 | 6975 | 1942.632 |
|  | 5 | RT20-5 | 78.71 | 161.40 | 3.78 | 3.33 | 12506 | 4705 | 1964.442 |
| RT21 | 1 | RT21-1 | 68.17 | 137.55 | 4.32 | 3.53 | 5371 | 6159 | 1933.004 |
|  | 2 | RT21-2 | 74.92 | 152.65 | 2.97 | 2.41 | 8964 | 4544 | 1951.931 |
|  | 3 | RT21-3 | 77.28 | 154.04 | 4.44 | 3.45 | 14982 | 6721 | 1909.554 |
|  | 4 | RT21-4 | 76.45 | 153.99 | 4.10 | 3.78 | 7373 | 6747 | 1929.659 |
|  | 5 | RT21-5 | 67.34 | 133.32 | 4.01 | 3.89 | 7203 | 3831 | 1896.652 |
| RT22 | 1 | RT22-1 | 71.77 | 145.51 | 3.60 | 3.10 | 6265 | 10704 | 1942.296 |
|  | 2 | RT22-2 | 79.51 | 159.50 | 3.99 | 2.98 | 7438 | 6717 | 1921.783 |
|  | 3 | RT22-3 | 70.08 | 141.48 | 3.70 | 3.73 | 8207 | 5413 | 1934.045 |
|  | 4 | RT22-4 | 84.03 | 167.10 | 3.90 | 3.42 | 9985 | 8419 | 1905.055 |
|  | 5 | RT22-5 | 79.04 | 160.60 | 3.91 | 3.94 | 5251 | 8283 | 1946.544 |
| RT23 | 1 | RT23-1 | 79.24 | 159.81 | 3.82 | 3.46 | 7420 | 8734 | 1932.080 |
|  | 2 | RT23-2 | 78.79 | 162.45 | 3.93 | 2.51 | 5094 | 5320 | 1975.214 |
|  | 3 | RT23-3 | 86.72 | 175.71 | 2.94 | 2.78 | 9005 | 6373 | 1941.077 |
|  | 4 | RT23-4 | 64.34 | 128.55 | 3.80 | 3.82 | 4535 | 5319 | 1914.064 |
|  | 5 | RT23-5 | 71.70 | 143.62 | 3.73 | 4.04 | 9000 | 4547 | 1918.939 |
| RT24 | 1 | RT24-1 | 89.73 | 179.43 | 2.91 | 2.74 | 10982 | 3578 | 1915.680 |
|  | 2 | RT24-2 | 69.34 | 140.31 | 3.71 | 3.39 | 12272 | 8091 | 1938.520 |
|  | 3 | RT24-3 | 74.88 | 151.10 | 3.27 | 3.75 | 8701 | 6076 | 1933.144 |
|  | 4 | RT24-4 | 68.25 | 135.08 | 4.60 | 3.26 | 5911 | 5514 | 1896.068 |
|  | 5 | RT24-5 | 68.71 | 139.42 | 4.33 | 3.25 | 4622 | 6055 | 1943.885 |
| RT25 | 1 | RT25-1 | 66.72 | 137.12 | 3.78 | 3.40 | 13317 | 6773 | 1968.839 |
|  | 2 | RT25-2 | 75.75 | 151.17 | 3.27 | 2.89 | 9169 | 7212 | 1911.827 |
|  | 3 | RT25-3 | 74.49 | 151.66 | 3.37 | 3.46 | 6444 | 6621 | 1950.467 |
|  | 4 | RT25-4 | 77.78 | 155.50 | 3.53 | 3.38 | 9779 | 5757 | 1915.261 |
|  | 5 | RT25-5 | 73.27 | 148.58 | 3.37 | 3.24 | 2840 | 4596 | 1942.673 |
| RT26 | 1 | RT26-1 | 80.29 | 162.97 | 3.50 | 3.09 | 4729 | 4565 | 1944.517 |
|  | 2 | RT26-2 | 67.33 | 136.19 | 3.86 | 3.59 | 8043 | 5412 | 1937.769 |
|  | 3 | RT26-3 | 79.50 | 154.35 | 3.57 | 3.51 | 14451 | 5300 | 1859.966 |
|  | 4 | RT26-4 | 78.46 | 158.37 | 3.92 | 3.20 | 10408 | 5879 | 1933.705 |
|  | 5 | RT26-5 | 66.21 | 128.52 | 4.09 | 3.93 | 4349 | 4335 | 1859.570 |
| RT27 | 1 | RT27-1 | 73.83 | 143.49 | 3.03 | 3.10 | 10179 | 5336 | 1861.891 |
|  | 2 | RT27-2 | 66.92 | 136.57 | 2.95 | 2.96 | 8397 | 4455 | 1955.082 |
|  | 3 | RT27-3 | 73.24 | 147.62 | 3.34 | 3.22 | 5518 | 6474 | 1930.912 |
|  | 4 | RT27-4 | 74.89 | 150.06 | 3.69 | 3.47 | 14038 | 7450 | 1919.582 |
|  | 5 | RT27-5 | 72.65 | 149.84 | 3.61 | 3.35 | 5300 | 4313 | 1975.867 |
| RT28 | 1 | RT28-1 | 83.20 | 166.45 | 4.71 | 3.74 | 25240 | 5188 | 1916.576 |
|  | 2 | RT28-2 | 69.45 | 142.65 | 3.63 | 3.48 | 6671 | 6258 | 1967.728 |
|  | 3 | RT28-3 | 73.50 | 150.98 | 3.52 | 3.65 | 4650 | 7316 | 1967.875 |
|  | 4 | RT28-4 | 72.17 | 148.06 | 3.74 | 3.03 | 8567 | 5799 | 1965.380 |
|  | 5 | RT28-5 | 66.28 | 132.90 | 3.39 | 2.73 | 7107 | 5880 | 1920.914 |
| RT29 | 1 | RT29-1 | 74.85 | 149.95 | 3.86 | 3.18 | 6270 | 7370 | 1919.200 |
|  | 2 | RT29-2 | 75.97 | 152.55 | 3.94 | 3.39 | 5234 | 5334 | 1923.692 |
|  | 3 | RT29-3 | 79.27 | 156.77 | 3.71 | 3.48 | 11138 | 6727 | 1894.609 |
|  | 4 | RT29-4 | 67.62 | 137.16 | 4.11 | 2.93 | 3510 | 5687 | 1943.201 |
|  | 5 | RT29-5 | 75.70 | 148.94 | 4.31 | 2.50 | 2816 | 5682 | 1884.868 |
| RT31 | 1 | RT31-1 | 70.45 | 144.73 | 3.51 | 2.93 | 5910 | 5514 | 1968.081 |
|  | 2 | RT31-2 | 78.61 | 163.09 | 3.95 | 3.18 | 3434 | 7109 | 1987.536 |
|  | 3 | RT31-3 | 84.63 | 169.11 | 4.06 | 3.37 | 9911 | 6169 | 1914.302 |
|  | 4 | RT31-4 | 73.36 | 146.21 | 3.78 | 3.22 | 1667 | 3187 | 1909.340 |
|  | 5 | RT31-5 | 67.25 | 134.85 | 3.62 | 3.61 | 2687 | 5598 | 1920.986 |
| RT32 | 1 | RT32-1 | 63.15 | 132.28 | 3.66 | 3.45 | 5071 | 6024 | 2006.718 |
|  | 2 | RT32-2 | 76.17 | 158.29 | 4.07 | 3.46 | 7643 | 5562 | 1990.834 |
|  | 3 | RT32-3 | 78.32 | 162.69 | 3.57 | 3.41 | 5660 | 5298 | 1990.003 |
|  | 4 | RT32-4 | 71.37 | 145.40 | 4.22 | 3.65 | 2973 | 6520 | 1951.705 |
|  | 5 | RT32-5 | 74.44 | 150.90 | 3.95 | 3.85 | 7126 | 5386 | 1941.996 |
| RT33 | 1 | RT33-1 | 69.75 | 143.83 | 3.45 | 3.24 | 7266 | 8203 | 1975.472 |
|  | 2 | RT33-2 | 72.92 | 146.97 | 3.77 | 2.85 | 2971 | 5895 | 1930.846 |
|  | 3 | RT33-3 | 84.38 | 168.77 | 4.38 | 3.38 | 9475 | 5431 | 1916.114 |
|  | 4 | RT33-4 | 76.02 | 154.34 | 3.99 | 3.18 | 3866 | 6186 | 1944.984 |
|  | 5 | RT33-5 | 68.97 | 141.91 | 3.78 | 3.61 | 5113 | 4535 | 1971.144 |
| RT34 | 1 | RT34-1 | 66.44 | 137.31 | 4.75 | 3.51 | 4926 | 4044 | 1979.876 |
|  | 2 | RT34-2 | 72.84 | 146.07 | 4.16 | 3.68 | 5057 | 4552 | 1921.129 |
|  | 3 | RT34-3 | 85.23 | 171.37 | 3.81 | 3.41 | 7108 | 10194 | 1926.229 |
|  | 4 | RT34-4 | 83.58 | 171.14 | 3.94 | 2.83 | 3773 | 6610 | 1961.619 |
|  | 5 | RT34-5 | 75.24 | 153.38 | 4.19 | 3.78 | 7329 | 5894 | 1952.925 |
| RT35 | 1 | RT35-1 | 80.84 | 165.93 | 3.57 | 3.66 | 18101 | 4664 | 1966.365 |
|  | 2 | RT35-2 | 62.10 | 125.74 | 4.08 | 3.22 | 15373 | 7159 | 1939.757 |
|  | 3 | RT35-3 | 59.02 | 119.68 | 4.44 | 4.62 | 9726 | 5006 | 1942.620 |
|  | 4 | RT35-4 | 68.07 | 138.69 | 3.42 | 2.87 | 4685 | 5717 | 1951.888 |
|  | 5 | RT35-5 | 65.71 | 134.18 | 4.38 | 3.52 | 4940 | 5467 | 1956.239 |
| RT36 | 1 | RT36-1 | 52.14 | 106.04 | 4.82 | 4.11 | 13832 | 4570 | 1948.338 |
|  | 2 | RT36-2 | 69.20 | 139.31 | 3.90 | 3.55 | 8595 | 5795 | 1928.598 |
|  | 3 | RT36-3 | 66.98 | 134.70 | 3.85 | 3.60 | 6206 | 2756 | 1926.584 |
|  | 4 | RT36-4 | 59.14 | 118.81 | 4.69 | 4.86 | 10786 | 2779 | 1924.585 |
|  | 5 | RT36-5 | 56.70 | 117.81 | 3.04 | 3.07 | 3220 | 4197 | 1990.511 |
| RT37 | 1 | RT37-1 | 68.40 | 138.60 | 3.22 | 3.91 | 7008 | 4749 | 1941.211 |
|  | 2 | RT37-2 | 80.46 | 166.54 | 3.71 | 3.02 | 7591 | 5769 | 1982.915 |
|  | 3 | RT37-3 | 64.02 | 131.28 | 4.29 | 3.85 | 5336 | 5012 | 1964.484 |
|  | 4 | RT37-4 | 71.40 | 143.73 | 4.27 | 4.18 | 7749 | 3914 | 1928.478 |
|  | 5 | RT37-5 | 69.41 | 139.90 | 4.04 | 3.89 | 5084 | 5823 | 1930.906 |
| RT38 | 1 | RT38-1 | 72.39 | 151.80 | 3.92 | 3.53 | 14416 | 5590 | 2008.902 |
|  | 2 | RT38-2 | 71.95 | 149.24 | 3.05 | 2.78 | 2980 | 4505 | 1987.101 |
|  | 3 | RT38-3 | 75.16 | 159.18 | 3.68 | 3.09 | 4513 | 5914 | 2028.931 |
|  | 4 | RT38-4 | 60.02 | 125.56 | 3.69 | 3.96 | 7688 | 4194 | 2004.107 |
|  | 5 | RT38-5 | 75.66 | 157.21 | 3.92 | 3.78 | 6049 | 7259 | 1990.579 |
| RT39 | 1 | RT39-1 | 75.57 | 154.09 | 3.69 | 3.82 | 6403 | 5196 | 1953.397 |
|  | 2 | RT39-2 | 69.26 | 144.51 | 4.14 | 3.13 | 2185 | 2570 | 1998.853 |
|  | 3 | RT39-3 | 72.51 | 148.03 | 3.24 | 3.85 | 6761 | 4891 | 1955.768 |
|  | 4 | RT39-4 | 72.97 | 149.22 | 4.40 | 3.24 | 2900 | 6506 | 1959.062 |
|  | 5 | RT39-5 | 64.10 | 130.35 | 3.73 | 3.34 | 4729 | 1967 | 1948.133 |
| RT40 | 1 | RT40-1 | 81.60 | 167.88 | 3.84 | 3.16 | 7309 | 5824 | 1970.944 |
|  | 2 | RT40-2 | 77.44 | 160.58 | 3.72 | 3.31 | 6084 | 6123 | 1986.514 |
|  | 3 | RT40-3 | 74.84 | 154.79 | 4.24 | 3.63 | 6208 | 6214 | 1981.411 |
|  | 4 | RT40-4 | 63.18 | 132.96 | 3.95 | 3.82 | 3536 | 5382 | 2016.076 |
|  | 5 | RT40-5 | 76.25 | 158.62 | 3.94 | 2.77 | 11671 | 5592 | 1992.891 |
| RT41 | 1 | RT41-1 | 77.23 | 159.12 | 3.13 | 3.29 | 5583 | 6536 | 1973.805 |
|  | 2 | RT41-2 | 77.77 | 160.15 | 3.73 | 3.21 | 5742 | 6904 | 1972.788 |
|  | 3 | RT41-3 | 80.75 | 172.14 | 3.38 | 3.57 | 6405 | 7749 | 2042.231 |
|  | 4 | RT41-4 | 76.91 | 157.58 | 3.79 | 3.70 | 3964 | 6783 | 1962.835 |
|  | 5 | RT41-5 | 69.21 | 146.01 | 4.05 | 3.68 | 2916 | 5064 | 2021.060 |
| RT42 | 1 | RT42-1 | 76.71 | 157.40 | 4.52 | 3.63 | 4866 | 5413 | 1965.705 |
|  | 2 | RT42-2 | 67.07 | 135.91 | 3.41 | 3.38 | 7208 | 5983 | 1941.282 |
|  | 3 | RT42-3 | 63.98 | 127.81 | 4.59 | 4.28 | 7811 | 5115 | 1913.754 |
|  | 4 | RT42-4 | 65.41 | 131.31 | 4.85 | 3.62 | 9638 | 6555 | 1923.177 |
|  | 5 | RT42-5 | 83.50 | 168.24 | 3.64 | 3.09 | 5986 | 7586 | 1930.227 |
| RT43 | 1 | RT43-1 | 83.07 | 167.19 | 3.29 | 3.10 | 6933 | 5563 | 1928.109 |
|  | 2 | RT43-2 | 76.16 | 152.05 | 3.97 | 3.78 | 8044 | 5739 | 1912.604 |
|  | 3 | RT43-3 | 79.04 | 158.66 | 4.27 | 3.37 | 8288 | 8786 | 1923.030 |
|  | 4 | RT43-4 | 84.21 | 166.73 | 3.40 | 3.51 | 6734 | 5589 | 1896.774 |
|  | 5 | RT43-5 | 75.12 | 147.96 | 3.88 | 3.61 | 3549 | 5961 | 1886.923 |
| RT44 | 1 | RT44-1 | 71.15 | 148.18 | 4.55 | 4.05 | 5479 | 5305 | 1995.171 |
|  | 2 | RT44-2 | 66.16 | 137.53 | 4.70 | 3.55 | 3078 | 5279 | 1991.441 |
|  | 3 | RT44-3 | 71.76 | 147.56 | 3.54 | 3.56 | 7312 | 3099 | 1969.934 |
|  | 4 | RT44-4 | 79.95 | 163.89 | 3.72 | 3.23 | 3748 | 5125 | 1963.810 |
|  | 5 | RT44-5 | 85.42 | 172.07 | 3.91 | 3.97 | 9970 | 3696 | 1929.795 |
| RT45 | 1 | RT45-1 | 69.30 | 145.23 | 3.93 | 3.65 | 6883 | 7415 | 2007.653 |
|  | 2 | RT45-2 | 83.18 | 172.37 | 3.84 | 2.71 | 6623 | 11997 | 1985.218 |
|  | 3 | RT45-3 | 63.16 | 132.96 | 3.95 | 3.83 | 3536 | 5385 | 2016.714 |
|  | 4 | RT45-4 | 76.02 | 155.63 | 4.14 | 3.51 | 6154 | 6988 | 1961.241 |
|  | 5 | RT45-5 | 71.95 | 145.72 | 3.79 | 3.85 | 3528 | 5291 | 1940.233 |
| RT47 | 1 | RT47-1 | 69.79 | 144.06 | 3.56 | 2.86 | 3478 | 5704 | 1977.496 |
|  | 2 | RT47-2 | 68.97 | 138.33 | 4.79 | 3.60 | 4333 | 4707 | 1921.417 |
|  | 3 | RT47-3 | 82.43 | 169.00 | 4.41 | 3.77 | 5131 | 7499 | 1964.115 |
|  | 4 | RT47-4 | 74.21 | 150.87 | 3.53 | 3.37 | 5514 | 5998 | 1947.628 |
|  | 5 | RT47-5 | 81.75 | 168.79 | 3.88 | 3.52 | 3849 | 5956 | 1977.992 |
| RT48 | 1 | RT48-1 | 76.19 | 159.74 | 3.93 | 3.63 | 4865 | 5426 | 2008.543 |
|  | 2 | RT48-2 | 81.96 | 170.96 | 3.93 | 3.26 | 7945 | 6087 | 1998.288 |
|  | 3 | RT48-3 | 78.79 | 159.97 | 3.95 | 3.40 | 5455 | 5521 | 1945.060 |
|  | 4 | RT48-4 | 79.61 | 162.55 | 3.67 | 3.77 | 7190 | 6071 | 1956.072 |
|  | 5 | RT48-5 | 81.90 | 166.33 | 4.04 | 3.68 | 6680 | 5646 | 1945.594 |
| RT49 | 1 | RT49-1 | 73.30 | 153.43 | 3.35 | 2.95 | 10639 | 6787 | 2005.265 |
|  | 2 | RT49-2 | 69.27 | 145.09 | 3.71 | 3.87 | 3912 | 3180 | 2006.586 |
|  | 3 | RT49-3 | 80.81 | 165.66 | 4.45 | 3.84 | 3948 | 5893 | 1963.894 |
|  | 4 | RT49-4 | 75.16 | 156.18 | 3.80 | 3.68 | 7831 | 6020 | 1990.692 |
|  | 5 | RT49-5 | 79.71 | 166.40 | 4.08 | 3.25 | 4840 | 5687 | 1999.890 |
| RT50 | 1 | RT50-1 | 70.28 | 141.50 | 3.84 | 3.38 | 3965 | 6293 | 1928.813 |
|  | 2 | RT50-2 | 80.89 | 167.09 | 4.25 | 3.89 | 6248 | 5362 | 1978.888 |
|  | 3 | RT50-3 | 78.54 | 157.21 | 4.24 | 3.65 | 7413 | 6197 | 1917.586 |
|  | 4 | RT50-4 | 73.45 | 147.50 | 3.98 | 3.45 | 6432 | 5200 | 1923.826 |
|  | 5 | RT50-5 | 80.43 | 164.60 | 3.96 | 3.22 | 6322 | 5762 | 1960.547 |
| RT51 | 1 | RT51-1 | 77.36 | 154.78 | 3.82 | 3.49 | 5555 | 5295 | 1916.743 |
|  | 2 | RT51-2 | 62.89 | 129.99 | 3.99 | 3.56 | 5615 | 6815 | 1980.131 |
|  | 3 | RT51-3 | 81.00 | 161.98 | 3.80 | 3.63 | 9084 | 5067 | 1915.763 |
|  | 4 | RT51-4 | 76.75 | 156.08 | 3.08 | 2.83 | 6524 | 6090 | 1948.204 |
|  | 5 | RT51-5 | 73.68 | 149.59 | 4.09 | 3.27 | 6729 | 6802 | 1944.995 |
| RT52 | 1 | RT52-1 | 84.29 | 175.43 | 4.03 | 3.27 | 5165 | 2431 | 1993.854 |
|  | 2 | RT52-2 | 69.63 | 145.55 | 4.00 | 3.89 | 5019 | 5427 | 2002.541 |
|  | 3 | RT52-3 | 75.08 | 157.24 | 3.70 | 3.43 | 7185 | 6464 | 2006.339 |
|  | 4 | RT52-4 | 72.30 | 150.19 | 3.89 | 3.92 | 13433 | 6270 | 1990.069 |
|  | 5 | RT52-5 | 75.42 | 154.37 | 3.81 | 3.88 | 8379 | 7028 | 1960.839 |
| RT53 | 1 | RT53-1 | 80.17 | 160.82 | 3.86 | 3.56 | 5034 | 6266 | 1921.736 |
|  | 2 | RT53-2 | 77.62 | 158.16 | 2.92 | 2.85 | 5011 | 6083 | 1952.039 |
|  | 3 | RT53-3 | 83.54 | 171.01 | 3.68 | 3.59 | 6995 | 6735 | 1961.068 |
|  | 4 | RT53-4 | 72.72 | 148.43 | 3.81 | 3.85 | 2725 | 4888 | 1955.390 |
|  | 5 | RT53-5 | 72.95 | 148.33 | 3.27 | 3.98 | 14494 | 4318 | 1947.911 |
| RT54 | 1 | RT54-1 | 69.35 | 146.98 | 4.05 | 3.74 | 6271 | 9274 | 2030.380 |
|  | 2 | RT54-2 | 83.35 | 172.06 | 3.69 | 3.23 | 5341 | 5108 | 1977.606 |
|  | 3 | RT54-3 | 85.92 | 176.29 | 4.05 | 3.23 | 3759 | 5612 | 1965.617 |
|  | 4 | RT54-4 | 74.13 | 155.45 | 3.71 | 3.60 | 4157 | 6193 | 2008.918 |
|  | 5 | RT54-5 | 68.73 | 143.34 | 3.86 | 3.63 | 6204 | 7441 | 1997.959 |
| RT55 | 1 | RT55-1 | 80.75 | 158.35 | 3.60 | 3.50 | 6759 | 6756 | 1878.629 |
|  | 2 | RT55-2 | 64.12 | 127.60 | 4.29 | 3.77 | 6513 | 7091 | 1906.438 |
|  | 3 | RT55-3 | 74.75 | 148.98 | 3.75 | 3.00 | 7812 | 8832 | 1909.336 |
|  | 4 | RT55-4 | 67.72 | 135.19 | 4.76 | 3.71 | 3943 | 6816 | 1912.463 |
|  | 5 | RT55-5 | 76.26 | 147.75 | 4.73 | 3.47 | 6909 | 5814 | 1856.078 |
| RT56 | 1 | RT56-1 | 70.03 | 143.24 | 4.63 | 3.91 | 5253 | 4961 | 1959.502 |
|  | 2 | RT56-2 | 72.76 | 146.78 | 3.25 | 3.11 | 5425 | 3931 | 1932.590 |
|  | 3 | RT56-3 | 74.56 | 149.16 | 3.68 | 3.27 | 5510 | 5167 | 1916.514 |
|  | 4 | RT56-4 | 76.78 | 154.51 | 3.93 | 3.00 | 10045 | 6686 | 1927.853 |
|  | 5 | RT56-5 | 76.95 | 156.05 | 3.79 | 2.84 | 6904 | 3363 | 1942.767 |
