## Supplementary material for "The small genome size ensures adaptive flexibility for an alpine ginger": Table S3

| **Table S3** Morphological traits of wild and common-garden populations for *Roscoea tibetica*. | | | | | | | |
| --- | --- | --- | --- | --- | --- | --- | --- |
| Habitat | Populations | Altitude (m) | GS (Gb) | B.ASS | F.ASS | B.ASN | F.ASN |
| Field | RT53 | 3023 | 1.94762873 | 51.08313108 | 51.15849185 | 73.05595238 | 4.131904762 |
| Field | RT54 | 2731 | 1.99609604 | 50.69311098 | 50.61243564 | 65.18630952 | 1.29047619 |
| Field | RT50 | 3343 | 1.94193188 | 48.97601184 | 49.20976018 | 98.2978836 | 5.006349206 |
| Field | RT47 | 3160 | 1.95772963 | 46.78930788 | 47.00141561 | 99.9021164 | 7.607407407 |
| Field | RT48 | 2602 | 1.97071142 | 45.76880115 | 45.58256058 | 146.8309524 | 12.03630952 |
| Field | RT52 | 3123 | 1.99072829 | 49.23030997 | 49.47914309 | 89.38888889 | 4.294444444 |
| Field | RT8 | 2888 | 1.9720285 | 48.15748506 | 48.24713814 | 112.9071429 | 6.616666667 |
| Field | RT20 | 3634 | 1.94975895 | 41.77940313 | 41.8409849 | 130.2797619 | 8.270833333 |
| Field | RT11 | 2898 | 1.92782046 | 43.34741986 | 43.34136708 | 156.65 | 8.2375 |
| Field | RT10 | 3028 | 1.94598782 | 39.18021926 | 39.11347569 | 164.6634524 | 12.65793651 |
| Field | RT1 | 2555 | 1.9870753 | 49.4783816 | 48.93330572 | 96.65185185 | 3.762962963 |
| Field | RT20 | 3634 | 1.94975895 | 45.27507747 | 45.25439319 | 110.4363095 | 4.64047619 |
| Field | RT21 | 3333 | 1.92416011 | 49.44718311 | 49.39681868 | 83.025 | 4.5995671 |
| Field | RT19 | 3451 | 1.926911 | 42.65372079 | 42.92538927 | 121.6 | 7.100529101 |
| Field | RT15 | 3263.5 | 1.92158295 | 44.1867113 | 44.75085807 | 91.37142857 | 5.321428571 |
| Field | RT38 | 2979 | 2.00392378 | 46.99286839 | 47.20427495 | 50.53401361 | 4.433333333 |
| Field | RT25 | 3742 | 1.93781332 | 42.73736293 | 42.48427612 | 128.8238095 | 10.87797619 |
| Field | RT14 | 2799 | 1.95010381 | 46.03813162 | 46.14245234 | 107.6190476 | 7.032407407 |
| Field | RT16 | 3249 | 1.96006705 | 51.9976528 | 51.74709174 | 66.61632653 | 3.980952381 |
| Field | RT18 | 3569 | 1.928273 | 44.84187202 | 44.32580855 | 109.8390476 | 3.373809524 |
| Field | RT7 | 3219 | 1.99531872 | 50.51749602 | 50.55205955 | 63.22592593 | 3.059259259 |
| Field | RT6 | 2569 | 2.00167974 | 49.36587412 | 49.23023101 | 93.88 | 4.226666667 |
| Field | RT12 | 2802 | 1.94538626 | 46.70524085 | 46.46128299 | 106.1886905 | 3.435714286 |
| Common garden | RT51 |  | 1.94116716 | 48.64233049 | 48.70753806 | 85.76666667 | 2.774285714 |
| Common garden | RT53 |  | 1.94762873 | 49.885209 | 49.84752337 | 56.95 | 2.156190476 |
| Common garden | RT54 |  | 1.99609604 | 50.2227274 | 49.94515362 | 42.95 | 0.776666667 |
| Common garden | RT50 |  | 1.94193188 | 51.61682675 | 51.07157393 | 76.99333333 | 3.758095238 |
| Common garden | RT47 |  | 1.95772963 | 45.05069058 | 45.49552353 | 98.06666667 | 6.847619048 |
| Common garden | RT8 |  | 1.9720285 | 49.27573407 | 49.02005564 | 56.77248677 | 1.769791667 |
| Common garden | RT11 |  | 1.92782046 | 48.16170669 | 48.19422845 | 67.12666667 | 1.086666667 |
| Common garden | RT20 |  | 1.94975895 | 48.52588267 | 48.49932293 | 96.63333333 | 2.085714286 |
| Common garden | RT21 |  | 1.92416011 | 51.39013483 | 51.51792417 | 54.81333333 | 2.616666667 |
| Common garden | RT19 |  | 1.926911 | 48.12060758 | 47.96190813 | 84.22 | 2.576190476 |
| Common garden | RT15 |  | 1.92158295 | 59.73143303 | 59.37377464 | 63.38095238 | 1.401360544 |
| Common garden | RT38 |  | 2.00392378 | 48.18188351 | 48.19217555 | 65.695 | 2.803333333 |
| Common garden | RT25 |  | 1.93781332 | 46.44643242 | 46.7482079 | 78.64666667 | 5.045714286 |
| Common garden | RT16 |  | 1.96006705 | 50.98873884 | 50.84111787 | 62.30555556 | 2.994444444 |
| Common garden | RT18 |  | 1.928273 | 47.72833026 | 47.36817976 | 82.75 | 2.28 |
| Common garden | RT7 |  | 1.99531872 | 48.31634005 | 48.12383508 | 66.44666667 | 2.144285714 |
| Common garden | RT6 |  | 2.00167974 | 51.31056498 | 51.47980057 | 70 | 3.128571429 |
| R, *Roscoea tibetica*; GS, population-average genome size estimates of 5 individuals. | | | | |  |  |  |
| B.ASS, the average stomatal size on the abaxial of leaf; F.ASS, the average stomatal size on the adaxial of leaf. | | | | | | |  |
| B.ASN, the average stomatal number on the abaxial of leaf; F.ASN, the average stomatal number on the adaxial of leaf. | | | | | | |  |
