## Supplementary material for "The small genome size ensures adaptive flexibility for an alpine ginger": Table S4

| **Table S4** Genome size variations between days for *Roscoea tibetica*. | | | | | | | | | | | | | |
| --- | --- | --- | --- | --- | --- | --- | --- | --- | --- | --- | --- | --- | --- |
| **Individual** | **Day** | **Sample numbers** | **Gain** | **Mean-X** | | **CV-X%** | | **Count** | | **C-value/ Mb** | **SD** | **Average C-value/ Mb** | **CV between days %** |
|  |  |  |  | **Tomato** | ***R. tibetica*** | **Tomato** | ***R. tibetica*** | **Tomato** | ***R. tibetica*** |  |  |  |  |
| RT2 | 1 | RT2-1 | 500 | 28.44 | 59.54 | 3.60 | 3.17 | 4165 | 8128 | 2005.602 | 19.097 | 1983.872 | 0.963 |
|  | 2 | RT2-2 | 500 | 28.46 | 58.71 | 3.61 | 3.21 | 3484 | 8472 | 1976.254 |  |  |  |
|  | 3 | RT2-3 | 500 | 28.69 | 58.99 | 4.63 | 3.70 | 4901 | 5106 | 1969.760 |  |  |  |
| RT10 | 1 | RT10-1 | 500 | 28.62 | 58.22 | 3.52 | 2.83 | 5564 | 7787 | 1948.804 | 15.392 | 1933.466 | 0.796 |
|  | 2 | RT10-2 | 500 | 23.72 | 47.49 | 4.92 | 3.74 | 3742 | 7286 | 1918.019 |  |  |  |
|  | 3 | RT10-3 | 500 | 26.71 | 53.91 | 3.64 | 2.74 | 4030 | 3530 | 1933.575 |  |  |  |
| RT16 | 1 | RT16-1 | 500 | 27.83 | 56.73 | 4.00 | 3.15 | 4363 | 4453 | 1952.833 | 16.073 | 1935.018 | 0.831 |
|  | 2 | RT16-2 | 500 | 27.35 | 54.86 | 4.37 | 2.96 | 3379 | 7499 | 1921.604 |  |  |  |
|  | 3 | RT16-3 | 500 | 28.18 | 56.79 | 3.77 | 3.21 | 4260 | 1832 | 1930.618 |  |  |  |
| RT20 | 1 | RT16-1 | 500 | 25.47 | 51.08 | 4.21 | 3.72 | 5468 | 4949 | 1921.266 | 23.619 | 1919.892 | 1.230 |
|  | 2 | RT16-2 | 500 | 23.03 | 45.57 | 5.44 | 3.30 | 6562 | 8103 | 1895.617 |  |  |  |
|  | 3 | RT16-3 | 500 | 21.81 | 44.23 | 4.46 | 3.52 | 4248 | 4892 | 1942.794 |  |  |  |
| RT24 | 1 | RT24-1 | 500 | 24.62 | 49.44 | 4.12 | 3.68 | 5068 | 4232 | 1923.782 | 22.137 | 1902.587 | 1.164 |
|  | 2 | RT24-2 | 500 | 23.17 | 45.46 | 4.24 | 3.21 | 4836 | 15839 | 1879.615 |  |  |  |
|  | 3 | RT24-3 | 500 | 24.70 | 49.10 | 4.35 | 3.90 | 5665 | 7710 | 1904.364 |  |  |  |
| RT25 | 1 | RT25-1 | 500 | 25.79 | 51.89 | 4.25 | 3.96 | 5116 | 3417 | 1927.515 | 9.959 | 1916.071 | 0.520 |
|  | 2 | RT25-2 | 500 | 24.59 | 49.01 | 4.75 | 3.44 | 5160 | 6110 | 1909.377 |  |  |  |
|  | 3 | RT25-3 | 500 | 24.57 | 49.02 | 4.11 | 3.21 | 4390 | 2211 | 1911.321 |  |  |  |
| RT26 | 1 | RT26-1 | 500 | 26.05 | 51.93 | 4.19 | 3.73 | 5047 | 6255 | 1909.748 | 27.303 | 1889.305 | 1.445 |
|  | 2 | RT26-2 | 500 | 25.07 | 48.63 | 5.62 | 3.63 | 5170 | 3645 | 1858.298 |  |  |  |
|  | 3 | RT26-3 | 500 | 27.91 | 55.35 | 3.69 | 2.90 | 9556 | 4775 | 1899.867 |  |  |  |
| RT28 | 1 | RT28-1 | 500 | 23.46 | 45.53 | 4.81 | 4.03 | 4440 | 4714 | 1859.239 | 24.321 | 1846.067 | 1.317 |
|  | 2 | RT28-2 | 500 | 23.15 | 44.97 | 4.44 | 3.63 | 4374 | 3801 | 1860.962 |  |  |  |
|  | 3 | RT28-3 | 500 | 25.71 | 48.79 | 4.07 | 3.15 | 4222 | 4500 | 1818.002 |  |  |  |
| RT42 | 1 | RT42-1 | 500 | 24.30 | 49.56 | 4.77 | 3.73 | 4987 | 5177 | 1953.847 | 6.081 | 1955.945 | 0.311 |
|  | 2 | RT42-2 | 500 | 24.77 | 50.75 | 4.42 | 3.47 | 5435 | 4267 | 1962.798 |  |  |  |
|  | 3 | RT42-3 | 500 | 24.50 | 49.90 | 4.02 | 3.17 | 3989 | 4864 | 1951.192 |  |  |  |
| RT55 | 1 | RT55-1 | 500 | 25.04 | 48.67 | 5.00 | 3.76 | 5044 | 9485 | 1862.055 | 3.286 | 1865.045 | 0.176 |
|  | 2 | RT55-2 | 500 | 24.84 | 48.45 | 4.51 | 3.64 | 4352 | 5138 | 1868.563 |  |  |  |
|  | 3 | RT55-3 | 500 | 26.61 | 51.79 | 3.62 | 2.82 | 3766 | 5112 | 1864.518 |  |  |  |
