## Supplementary Figure S1-S3 for "The small genome size ensures adaptive flexibility for an alpine ginger"

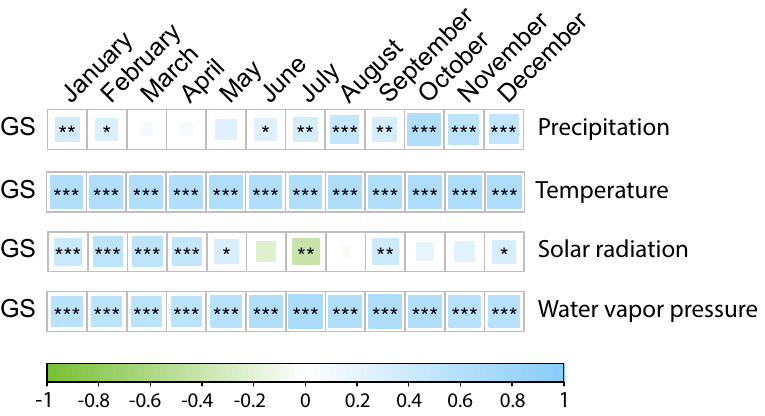

**Fig. S1** Pearson correlation coefficients between intraspecific genome size (GS) and monthly values of precipitation, temperature, solar radiation and water vapor pressure. Blue indicates positive correlation, green indicates negative correlation, and the darker the color, the stronger the correlation. Significance levels are denoted by asterisks: ^*^*p* < 0.05; ^**^*p* < 0.01; ^***^*p* < 0.001. The asterisk not shown means that the correlation is not significant.

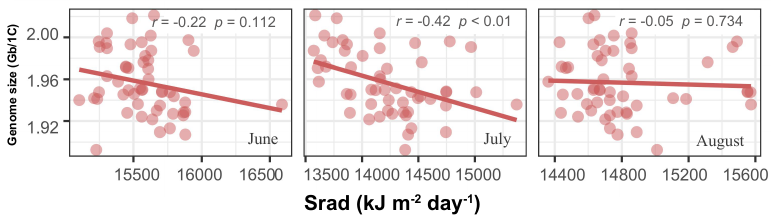

**Fig. S2** Scatter plot showing the correlation between genome size and solar radiation (Srad) of growth season in June, July and August for *Roscoea tibetica*. The solid red line represents the best fit linear regression.

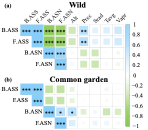

**Fig. S3** Correlations between morphological traits and environmental factors for the *Roscoea tibetica* from the wild **(a)** and the common garden **(b)**. Blue indicates positive correlation, green indicates negative correlation, and the darker the color, the stronger the correlation. Significance levels are denoted by asterisks: ^*^*p* < 0.05; ^**^*p* < 0.01; ^***^*p* < 0.001. The asterisk not shown means that the correlation is not significant. Abbreviations: B.ASS, the average stomatal size on the abaxial of leaf; F.ASS, the average stomatal size on the adaxial of leaf; B.ASN, the average stomatal number on the abaxial of leaf; F.ASN, the average stomatal number on the adaxial of leaf; Alt, altitude; Long, longitude; Lat, latitude; Prec, precipitation; Srad, solar radiation; Tavg, monthly average temperature; Vapr, water vapor pressure.
